# The sex determination gene *fruitless* is essential for male *Aedes aegypti* mosquito swarming and attraction to female flight sounds

**DOI:** 10.64898/2026.02.05.703956

**Authors:** YuMin M Loh, Alberto J Thornton, Tai-Ting Lee, Wan-Tze Chen, Yifeng Y.J. Xu, Daniel F Eberl, Matthew P Su, Azusa Kamikouchi

**Author notes:** These authors contributed equally (Matthew P Su, Azusa Kamikouchi). Correspondence (M.P.S.), (A.K.).

## Abstract

Aggregating male mosquitoes locate females by showing strong and innate attraction to female flight sounds, with these behaviors forming the basis of mating in the world’s deadliest animal. In contrast, females do not show aggregation or sound attraction behaviors. Despite being documented for almost a century, the molecular underpinnings of these behavioral dimorphisms remain unknown. Here, we found that by disrupting the male-specific isoform of the key sex determination gene *fruitless* (referred to as *fruM*) in *Aedes aegypti*, mutant males behave like females. *fruM* mutant males show complete abolition of aggregation and sound attraction behaviors, resulting in failure to locate females for mating. We identified strong *fruitless* expression in control male ears, with the loss of FruM severely disrupting mutant male ability to amplify female flight sounds compared to control males. *fruM* mutant male ears propagate frequency-specific, sound-evoked electrical signals at about one-sixth the magnitude of control males, significantly reducing the amount of auditory information relayed to the brain. *fruitless* thus plays a significant role in determining proper male peripheral hearing function, which enables male hearing behaviors essential for mating. Transcriptomic analyses of mosquito ears enabled identification of essential hearing genes under the transcriptional influence of FruM. We propose that FruM fine-tunes the masculinization of mosquito ears through shaping male-specific hearing cellular machineries that support male attraction to female flight sounds. We conclude that FruM is the master regulator of male-specific reproductive behaviors in mosquitoes and propose *fruitless*, as well as *fruitless*-dependent genes, as novel targets for control strategies.

**Highlights:**

- FruM is the master regulator of male swarming and attraction to female flight sounds
- FruM in male ears is essential for male amplification of female flight sounds
- FruM in male ears modulates sound-evoked electrical signal propagation to the brain
- FruM regulates essential hearing gene expression

## Introduction

Sexual dimorphisms in insect behaviors arise from the need to divide reproductive tasks^1^, which in turn require corresponding dimorphisms in relevant sensory organs involved in the processing of sensory information^1,2^. In disease-transmitting mosquitoes, males form male-dominated mating swarms at dusk and show robust attraction to female flight sounds (‘phonotaxis’) to locate females^3,4^. Females neither swarm nor exhibit phonotaxis, but seek hosts and oviposit^3–7^.

Mosquito ears are composed of a flagellum coupled with a Johnston’s Organ (JO) housed in the second antennal segment, the pedicel^8^. Unlike other insect species, but like vertebrate ears, mosquito JOs are innervated by auditory efferents descending from the central nervous system (CNS)^9^. Sexual dimorphisms in mosquito ears are apparent at various levels: from the anatomical structure of the flagellum to the number and properties of JO neurons to the localization of auditory efferent terminals, these properties collectively shape dimorphisms in peripheral hearing function^10–12^. Understanding these anatomical and functional dimorphisms represents the first step towards understanding sex-specific hearing behaviors in mosquitoes.

In insects, sex-specific somatic cell proliferation and neuronal differentiation are differentially regulated by *doublesex* (*dsx*) and *fruitless* (*fru*), respectively^13,14^. The former is crucial in shaping sex-specific morphological structures while the latter determines sexual dimorphisms in behaviors^15,16^. Both *dsx* and *fru* encode transcription factors (TFs) that exert regulatory function by influencing downstream gene expression programs^17,18^. *dsx* transcripts undergo sex-specific alternative splicing that give rise to sex-specific Dsx isoforms: DsxF and DsxM in females and males, respectively^19^. In contrast, the sex-specific alternative splicing of *fru* transcripts is predicted to only produce functional FruM in males^20,21^. The presence of an additional female-specific exon in female *fru* transcripts introduces a premature stop codon predicted to yield a truncated Fru protein^20,21^.

Prior work investigating FruM in *Aedes aegypti* (*Ae. aegypti*) mosquitoes found that *fruM* mutant males develop female-like host-seeking behavior, suggesting that FruM inhibits female behaviors in males^21^. Mutant males also exhibit mating failures and do not inseminate females, though the underlying causes remain unknown^21^. While the role of *dsx* in regulating mosquito ear anatomy and hearing function has previously been reported^22^, *fru*’s potential role remains entirely unexplored.

Here, we found that *fruM* mutant males completely lacked male swarming and phonotaxis behaviors, preventing mutant males from successful mating. Anatomical profiling confirmed strong, broad expression of *fru* in male JO neurons, suggesting *fru* could influence male hearing function, which underlies male hearing behaviors. Functionally, the loss of FruM dramatically altered several key hearing parameters in mutants compared to control males, including the complete abolition of male ear-unique self-sustained oscillations (SOs) that reflect male amplification of female flight sounds at 1000s-fold above baseline, as well as a strong reduction in auditory neural responses to frequency-specific stimuli.

Our anatomical and molecular profiling suggests that the altered hearing function of *fruM* males could arise from both changes in the efferent control of mutant ears and intrinsic changes in the FruM-dependent hearing cellular machineries. Through motif enrichment and protein-DNA structural analyses, we identified a putative FruM-binding DNA motif that allowed for *in-silico* validation of proposed transcriptional networks and identification of putative FruM direct binding genes, laying the groundwork for future investigation of FruM-dependent mechanisms that define sexual dimorphisms in mosquito peripheral hearing function.

We conclude that FruM is the master regulator of male aggregation and phonotaxis. FruM directs multilayered transcriptional networks to influence male-specific hearing function and behaviors, with hearing representing the primary sensory system involved in mosquito mating. We propose the mosquito peripheral auditory system as a new model for studying the sex determination pathway and highlight *fru* as a novel promising target for mosquito control.

## Results

### FruM mutation abolishes male aggregation and phonotaxis

Male *Ae. aegypti* mosquitoes aggregate at dusk to find mates^5,23^. Mating is initiated when males detect female flight sounds and rapidly approach females for mating, a hearing behavior known as “phonotaxis” (Figure 1A)^3^. Females neither swarm nor show phonotaxis^3^.

**Figure 1:**
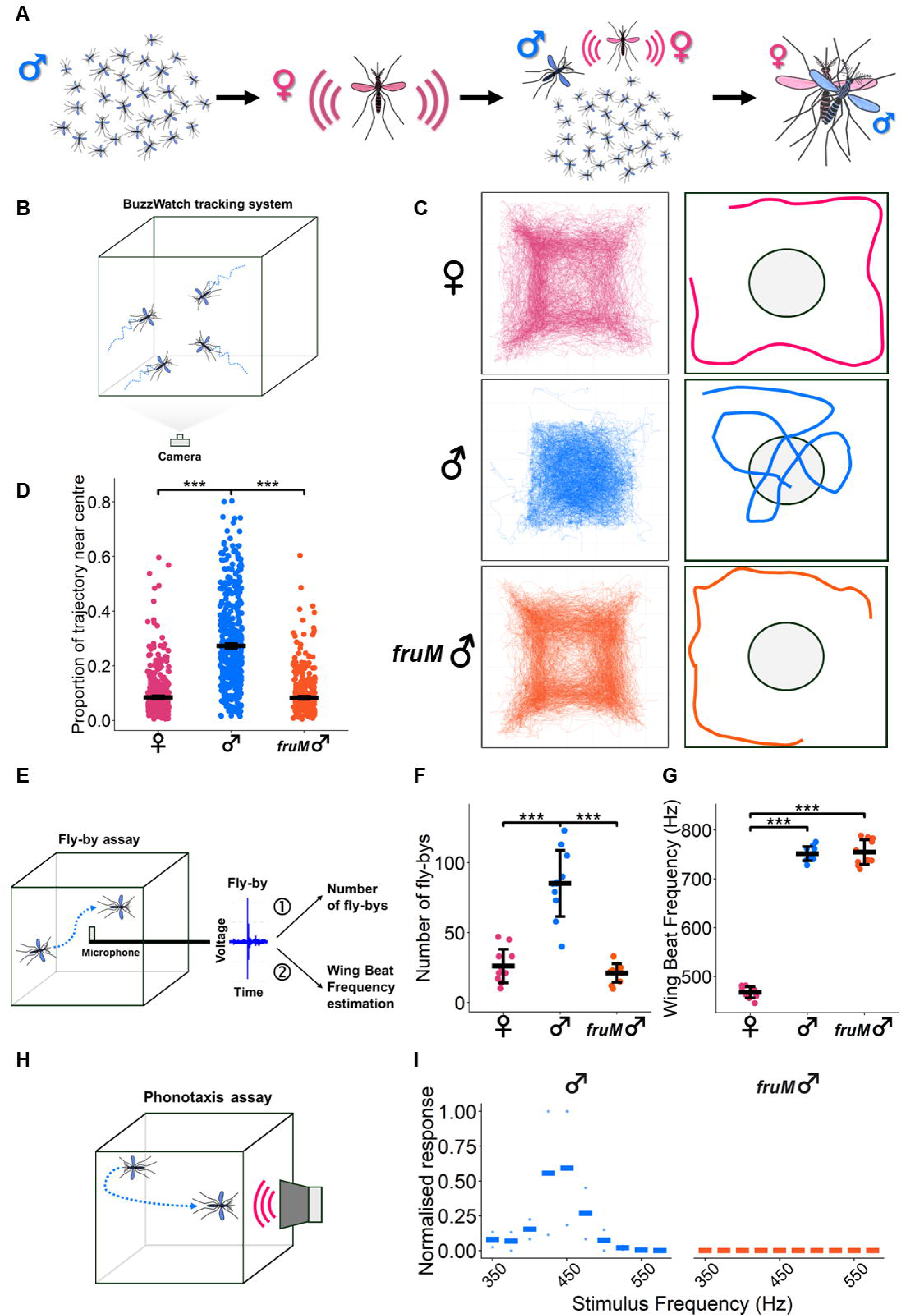
Loss of FruM abolishes male aggregation and phonotaxis. **(A)** Schematic of male aggregation and male attraction to female flight sounds. **(B)** Schematic illustration of BuzzWatch flight tracking system^24^. **(C)** Flight trajectories (left) from a cage of control female (top), control male (middle) and *fruM* male (bottom) mosquitoes between ZT11.5-12. 30 mosquitoes per cage. Schematic illustration of flight trajectories for each group tested, with circle of radius 100 pixels at centre (right, not to scale). **(D)** Quantification of proportion of mosquito flight trajectories within 100 pixels of cage centre. Individual points represent individual trajectories. Solid lines represent median and standard errors. See Table S3. ***, p<0.001, pairwise Wilcoxon tests with BH correction. Sample sizes: 250 control female trajectories; 392 control male trajectories; 232 *fruM* male trajectories. **(E)** WBF and fly-by assay diagram. **(F)** Number of fly-bys per cage for control female, control male and *fruM* male groups. Individual points represent total number of fly-bys for individual cages. Solid lines represent mean and standard deviations. See Tables 1 and S4. ***, p<0.001, pairwise t-tests with BH correction. Sample sizes: 11 control female cages; 11 control male cages; 11 *fruM* male cages. **(G)** Wing Beat Frequencies (WBFs) of control females, control males and *fruM* males. Individual points represent WBF estimates for one cage. Solid lines show mean and standard deviation. See Table S5. ***, p<0.001, pairwise t-test with BH correction. Sample sizes: 11 control females; 11 control males; 11 *fruM* males. **(H)** Phonotaxis assay diagram. **(I)** Normalised frequency response profiles of control males to a range of pure tones (350-575 Hz in 25 Hz bins). Individual points represent individual repeat normalised frequency responses. Bars represent median normalised response per frequency across all repeats. Sample sizes: 2 control male cages; 5 *fruM* male cages. See Figures S6A and S6B.

*fruM* mutant males exhibit mating failures, with unclear underlying causes^21^. To investigate this, we tested published heteroallelic *fruM* null mutant (*fruitless*^Δ*M*^*/fruitless*^Δ*M-tdTomato*^) males (hereafter denoted “*fruM* males”)^21^.

Since locomotion and flight are essential for mating, we first tested the baseline locomotor activity level of *fruM* males in a 12:12 Light:Dark cycle (Figure S1A). *fruM* males exhibited bimodal peaks in locomotor activity at dawn (ZT0) and dusk (ZT12), like control males and females (Figure S1B). However, compared to control males, *fruM* males showed a significant increase in locomotor activity in the morning (ZT2-4) and a significant decrease at dusk (ZT11-12) (pairwise Wilcoxon test with Benjamini Hochberg (BH) correction; p = 0.011 and p = 2.30×10^-5^; Tables S1 and S2), suggesting that FruM modulates male locomotion during specific time periods (Figures S1B and S1C).

Male swarming at dusk is essential for mating and strictly time-regulated^5^. The reduction in *fruM* male locomotor activity at dusk may reflect changes in swarming behavior. To investigate aggregation, we recorded cages of mosquitoes in the last 30 minutes before lights off to examine their flight patterns. We found that whilst control males tended to fly near the cage center, control females and *fruM* males typically flew along the cage sides (Extended data Video 1). We confirmed our observations using the BuzzWatch tracking system^24^ (Figures 1B, 1C and S2; Extended data Video 2). Significantly greater proportions of control male trajectories at dusk were within 100 pixels of the cage centre than control female and *fruM* male trajectories (pairwise Wilcoxon test with BH correction; p = 8.76 x 10^-47^ and p = 0; Figure 1D; Table S3), with heatmaps and individual trajectories providing further qualitative evidence of differences in flight patterns (Figures S2-S5).

We next placed a microphone into the center of a cage housing entrained, single-sex, free flying mosquitoes to record and extract: 1) the number of flyby events past the microphone, an indicator of male-aggregation at the cage center, and 2) mosquito Wing Beat Frequencies (WBFs) (Figure 1E). We found that the number of flybys for *fruM* males was significantly lower than control males but similar to control females (pairwise t-test with BH correction; p = 3.58 x 10^-6^ and p >0.05; Figure 1F; Tables 1 and S4). We found no significant difference in WBF between control and *fruM* males (pairwise t-test with BH correction; p >0.05), though control females had significantly lower WBFs than both male groups (p = 2.02 x 10^-21^ and p = 8.34 x 10^-15^; Figure 1G; Tables 1 and S5). We conclude that FruM controls male aggregation at dusk but does not influence male WBFs.

Since male attraction to female flight sounds is a prerequisite for courtship, *fruM* male mating failure could be due to primary defects in phonotaxis^21^. Using a phonotaxis assay to test male responses to a broad range of pure tones encompassing and extending beyond the female WBF range (Figure 1H), we found that *fruM* males failed to show attraction to all tones tested, while control and heterozygous *fruM* (*fru*^Δ*M*^/+) males were strongly attracted to specific tones (Figures 1I, S6A and 6B). Indeed, we found that even when directly presented with 470 Hz tones (matching the estimated control female WBF), only control males showed sound-induced abdominal bending behavior (Extended data Video 3).

We conclude that FruM is the master regulator of male *Ae. aegypti* reproductive behaviors, specifically male aggregation and phonotaxis.

### JO neurons are fru positive and FruM mutation alters male hearing function

Male peripheral hearing function underlies male phonotaxis^25^. We therefore hypothesized that the complete abolition of male phonotaxis in *fruM* males could be attributable to defects in mutant male peripheral hearing function.

We first examined *fru* expression in published wild-type male and female *Ae. aegypti* snRNA-seq and pedicel RNAseq databases^26,27^, and found strong and broad *fru* expression in the putative JO neuron cluster (i.e. auditory neuron cluster) (Figure 2A). We further confirmed that *fru* transcripts in the pedicel undergo sex-specific alternative splicing (Figure 2B)^21,27^. One of the two *fru* heteroallelic lines carries a tdTomato marker expressed under the control of the endogenous *fru* promoter, enabling visualization of *fru* expressing neurons^21^. We stained *fruitless*^Δ*M-tdTomato*^*/+* male JOs with anti-tdTomato and found extensive *fru* expression, consistent with snRNA-seq and pedicel RNAseq observations (Figures 2C and S7A).

**Figure 2:**
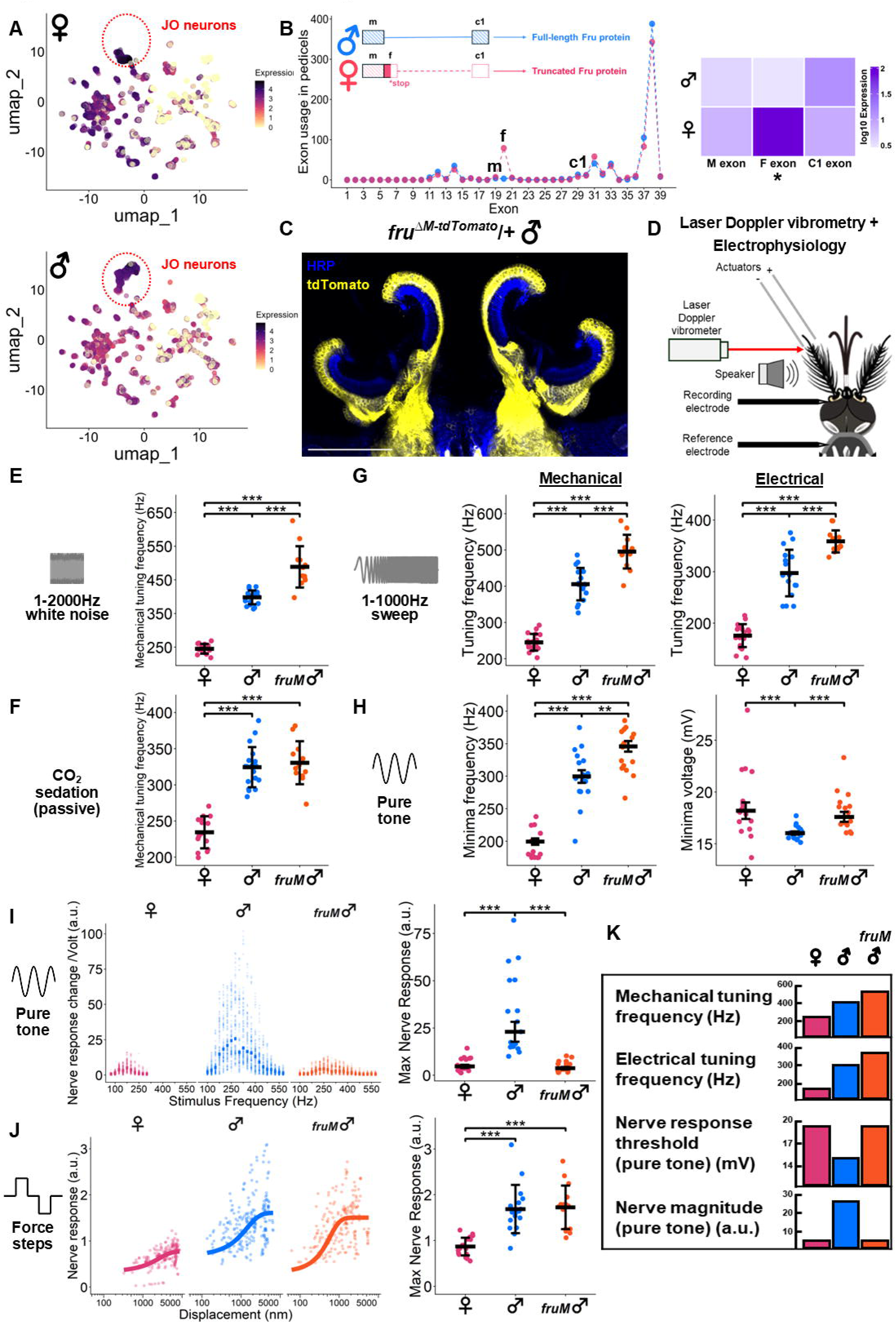
Loss of FruM alters male hearing function. **(A)** UMAP of head nuclei of male and female *Ae. aegypti* colored by normalized expression of *fru*. Data from^26^. Putative JO neurons, identified via expression of *inactive* (AAEL020482), are circled. **(B)** Schematic illustration of sex-specific alternative splicing of *fru* transcripts (left) and heatmap of exon usage counts of m, f and c1 exons in control female and control male pedicels as identified from reanalysis of published adult male and female *Ae. aegypti* pedicel transcriptome database^27^ (right). **(C)** Expression of *fru* in *fru*^Δ*M-tdTomato*^/+ male JO. Scale bar, 100 μm. Maximum intensity projection of JOs stained with anti-tdTomato (yellow) and anti-HRP (blue). See Figure S7A for split channel images. **(D)** Combined laser Doppler vibrometry and electrophysiology assay. **(E)** Active, white noise stimulated mechanical tuning frequencies of control females, control males and *fruM* males. Individual points represent measurements from individual mosquitoes. Solid lines represent means and standard deviations. ***, p<0.001, pairwise t-tests with BH correction. Sample sizes: 15 control females; 19 control males; 12 *fruM* males. **(F)** Passive, unstimulated mechanical tuning frequencies of control females, control males and *fruM* males. Individual points represent measurements from individual mosquitoes. Solid lines represent means and standard deviations. ***, p<0.001, pairwise t-tests with BH correction. Sample sizes: 15 control females; 19 control males; 12 *fruM* males. **(G)** Active, sweep stimulated mechanical (left) and electrical (right) tuning frequencies of control females, control males and *fruM* males. Individual points represent measurements from individual mosquitoes. Solid lines represent means and standard deviations. ***, p<0.001, pairwise t-tests with BH correction. Sample sizes: 19 control females; 17 control males; 13 *fruM* males. **(H)** Minima electrical tuning frequencies (left) and minima voltage threshold responses (right) from threshold pure tone stimulation experiments for control females, control males and *fruM* males. Individual points represent electrical tuning frequency estimates for individuals. Solid lines show median and standard error. **, p<0.01, ***, p<0.001, pairwise Wilcoxon tests with BH correction. Sample sizes: 17 control females; 17 control males; 16 *fruM* males. **(I)** Nerve phasic responses of control females, control males and *fruM* males from pure tone stimulation experiments. Individual points represent individual responses (nerve response magnitude normalized to stimulus voltage) to pure tones of the largest voltage intensity. Bars represent median responses to pure tones (left). Quantification of the maximum nerve phasic response for each individual tested (right). Sample sizes: 17 control females; 17 control males; 16 *fruM* males. **(J)** Nerve tonic responses of control females, control males and *fruM* males from force-step stimulation experiments. Individual points represent individual nerve responses to maximum flagellar displacements within an individual. Solid lines represent sigmoid fits (left). Quantification of the maximum nerve tonic response for each individual tested (right). Sample sizes: 17 control females; 17 control males; 15 *fruM* males. **(K)** Summary diagram of hearing parameters across genotypes.

Brain immunostaining of *fruitless*^Δ*M-tdTomato*^*/+* males showed strong tdTomato expression in the Antennal Mechanosensory and Motor Center (AMMC) to which JO neuron axons project^27,28^ (Figure S7B), confirming that JO neurons are *fru* positive. *Ae. aegypti* AMMCs show outstanding expression of *fru* relative to other neuropils, a greatly distinct pattern from that previously reported in fruit flies^29^. These observations suggest that FruM expression in male JOs may shape male-unique hearing function, thus determining mating success *via* regulating phonotaxis. Therefore, we next elucidated FruM’s influence on male peripheral hearing function.

Males use active hearing to detect female flight tones^30–32^. Mechanical deflections of the flagellum stretch-activates mechanotransducers in JO neurons to elicit action potentials that relay auditory information to the brain^33^. Mosquito hearing is a bidirectional process: in addition to mechanotransducing sound, JO neurons actively inject energy to set flagellae in motion even in the absence of sound^33^. This energy-dependent mechanical feedback amplification defines their baseline frequency sensitivity and selectivity for behaviorally relevant sounds^30–32^.

The free-fluctuating male flagellum actively vibrates at frequencies (mechanical tuning frequency) that encompass the range of female WBFs^33^, reflecting male ear active amplification of female flight sounds, which increases male hearing sensitivity and prepares males for when females enter his audible zone^34^. Frequency ranges of male and female hearing amplification appear different; male hearing amplification supports male phonotaxis, but the biological significance of female amplification is unclear^27^.

We first asked if FruM is involved in establishing male ear mechanical tuning frequency in response to sound, set by JO neuron motility. Mechanical vibrational properties of mosquito ears were measured *via* laser Doppler vibrometry (Figure 2D). We found significant differences between male and female stimulated mechanical tuning frequencies in response to broadband white-noise (pairwise t-test with BH correction; p = 3.36 x 10^-22^; Figure 2E; Tables 1 and S6). *fruM* males showed a significantly higher stimulated mechanical tuning frequency than control males (pairwise t-test with BH correction; 1.51 x 10^-5^; Figure 2E; Tables 1 and S6).

Active energy injection by JO neurons can be reversibly suppressed *via* CO_2_ sedation, enabling measurement of sex-specific differences in flagellar passive movements resulting from thermal bombardment, thus revealing structural and morphological differences of antennal ears between sexes^33^. Consistent with the lack of apparent anatomical differences between control and *fruM* male ears, passive ear mechanical tuning frequencies of male groups did not significantly differ from each other but were significantly different from control females (pairwise t-test with BH correction; p > 0.05 for male comparisons; p = 1.50 x 10^-12^ and p = 1.66 x 10^-8^; Figure 2F; Table S7). Our passive state measurements confirmed that active state mechanical tuning frequency differences between control and *fruM* males are determined by active JO neuron properties.

Regardless of the presence or absence of sound stimulation, control male active ear mechanical tuning frequency ranges encompassed the female WBF range, suggesting that control male ears tune to and actively amplify female flight sounds regardless of acoustic conditions (Figures 2E, S8A and S8B). However, we observed minimal overlap between *fruM* male ear mechanical tuning frequencies and female WBFs, revealing defects in mutant male ears in actively amplifying female flight sounds (Figure S8B).

After profiling ear mechanical vibrational properties, we next investigated ear electrical properties *via* electrophysiology (Figure 2D). We simultaneously measured electrical responses from the antennal nerve^25^, through which JO neurons relay signals to the brain, and ear mechanical vibrations. We started by quantifying responses to frequency sweep stimulation of 1-1000 Hz and determined ear electrical tuning frequency as the frequency eliciting the greatest compound action potential in the antennal nerve. We again found that *fruM* males had a significantly higher stimulated mechanical tuning frequency than control males (pairwise t-test with BH correction; p = 2.99 x 10^-4^; Figure 2G; Tables 2 and S8). Consistent with the mechanical tuning frequency phenotype, *fruM* male ear electrical tuning frequency was significantly higher than control males (pairwise t-test with BH correction; p = 4.79 x 10^-5^; Figure 2G; Tables 2 and S9).

To quantify changes in antennal nerve response magnitude upon FruM mutation, we determined the smallest stimulus magnitude necessary to induce a detectable antennal nerve response by providing pure tone stimuli (25 Hz bin) at increasing stimulus intensities^25^. Once again, the electrical tuning frequency of *fruM* males was significantly higher than control males and females (pairwise Wilcoxon tests with BH correction; p = 4.60 x 10^-3^ and p = 5.14 x 10^-9^; Figure 2H; Tables 2 and S10). *fruM* males had a significantly higher electrical response threshold compared to control males, but not control females (pairwise Wilcoxon tests with BH correction; p < 1.11 x 10^-4^ and p > 0.05; Figure 2H; Tables 2 and S11).

We observed that pure tone stimuli of identical intensities were able to elicit about six-fold greater antennal nerve response in control males compared to *fruM* males and control females (pairwise Wilcoxon tests with BH correction; p = 9.00 x 10^-9^ for both comparisons; Figure 2I; Table S12). Static deflections of ears elicited similar magnitudes of antennal nerve response in *fruM* and control males (pairwise t-tests with BH correction; p > 0.05; Figure 2J; Table S13).

The higher electrical response threshold and significantly reduced antennal nerve response of *fruM* male ears may reflect altered phasic responses of *fruM* male JO neurons to pure tone stimuli, potentially due to (feminized) alterations in mechanotransducer adaptation machinery^35–38^. The seemingly intact tonic response of *fruM* male JO neurons to static deflections suggests a similar number of functional mechanotransducer units between control and *fruM* males^33^.

Collectively, sex-specific alternative splicing of *fru* in JO neurons suggests that FruM regulates peripheral male auditory processing by establishing the sensitivity and selectivity of male ears to female flight sounds (Figure 2K).

### FruM influences localization of efferent presynaptic terminals in male JOs

We next characterized JO neuroanatomy *via* immunofluorescence, as sexual dimorphisms in hearing function are partly attributable to differences in JO neuroanatomy^33^.

We observed significant differences in JO width between control males and females (reflecting the substantially greater number of JO neurons in male ears ^28^), but not between control and *fruM* males, suggesting similar number of JO neurons in both male groups (pairwise t-test with BH correction; p < 4.36 x 10^-10^ and p > 0.05 respectively; Figure 3A; Tables S14 and S15), matching our functional characterization of their passive systems (Figure 2F).

**Figure 3:**
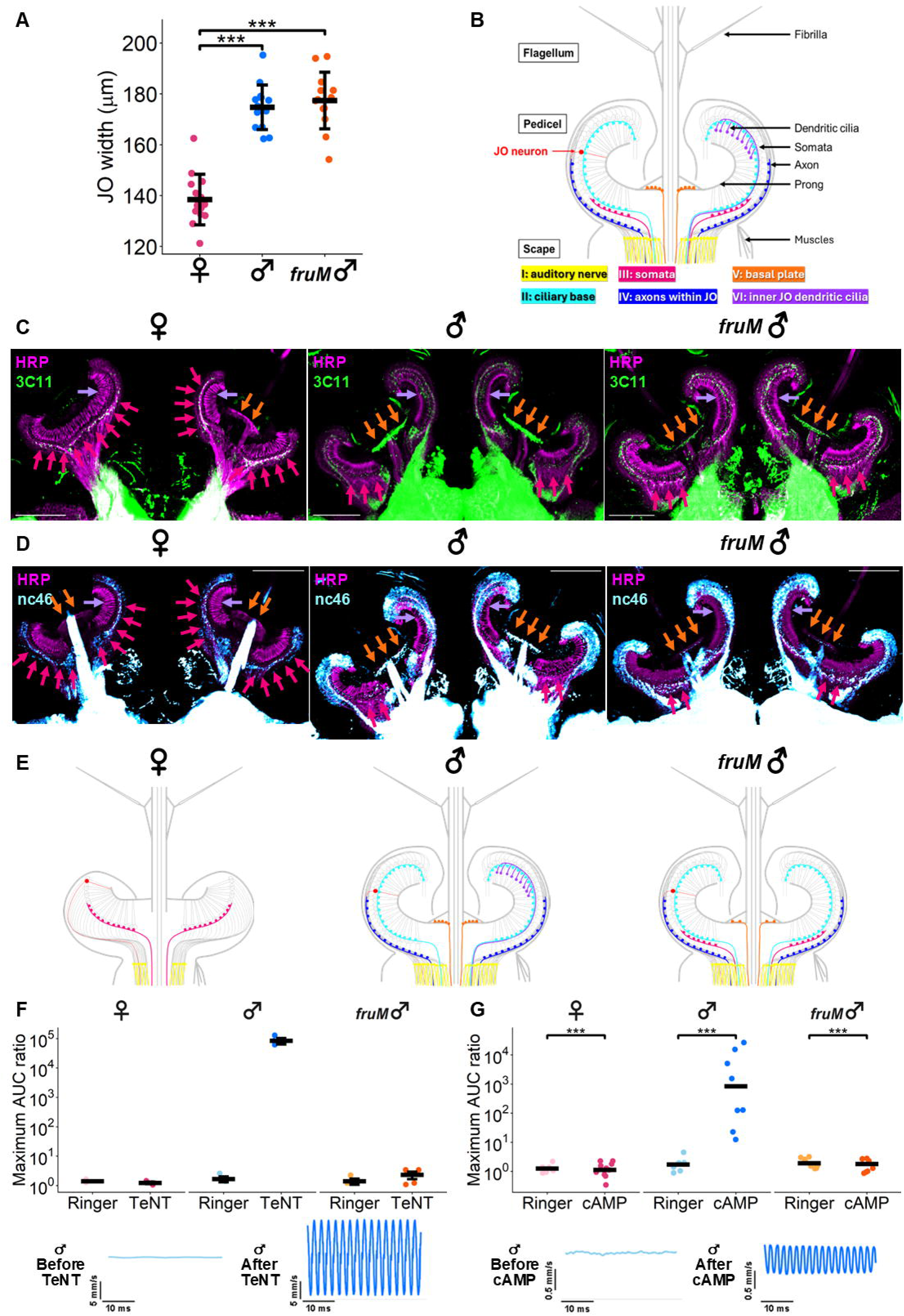
Loss of FruM alters localization of presynaptic terminals in male JOs and abolishes male SOs. **(A)** JO width of control females, control males, and *fruM* males. Individual points represent JO widths obtained from individual mosquito JOs. Solid lines represent mean and standard deviation. ***, p<0.001; pairwise t-tests with BH correction. Sample sizes: 14 control females; 14 control males; 13 *fruM* males. **(B)** Schematic of the different classes of efferent presynaptic terminal sites in mosquito JO. Adapted from Loh et al^10^. Type VI terminal represents a class first identified in this study. **(C)** Control female, control male and *fruM* male JO neuroanatomy visualized using immunofluorescence. Maximum intensity projections of JOs stained with anti-SYNORF1 (3C11, green) and anti-HRP (magenta). Scale bar, 100μm. Red arrows= type III terminals (somata), orange arrows= type V terminals (basal plate), purple arrows= type VI terminals (inner JO dendritic cilia). See Figure S9A for split channel images. **(D)** Control female, control male and *fruM* male JO neuroanatomy visualized using immunofluorescence. Maximum intensity projections of JOs stained with anti-SAP47 (nc46, blue) and anti-HRP (magenta). Scale bar, 100μm. Red arrows= type III terminals, orange arrows= type V terminals, purple arrows= type VI terminals. See Figure S9B for split channel images. **(E)** Summary schematic of presynaptic terminal locations across groups. Color coding of efferent terminal types follows Figure 3B. **(F)** Maximum ratio of AUC for vibrometry recordings taken before and after compound injection (Ringer or TeNT) for control females, control males and *fruM* males. Individual points represent data from individual mosquitoes. Solid lines show median and standard error. See Table S16 for maximum ratio of AUC following Ringer or TeNT injections (top). Representative control male flagellar vibrations before and after TeNT injection (bottom). Sample sizes for Ringer and TeNT injections: 3/3 control females; 3/3 control males; 3/4 *fruM* males. **(G)** Maximum ratio of AUC for vibrometry recordings taken before and after compound injection (Ringer or cAMP) for control females, control males and *fruM* males. Individual points represent data from individual mosquitoes. Solid lines show medians. ***, p < 0.001, Wilcoxon tests. See Table S17 for maximum ratio of AUC following Ringer or cAMP injections and p values (top). Representative control male flagellar vibrations before and after cAMP injection (bottom). Sample sizes for Ringer and cAMP injections: 7/10 control females; 8/8 control males; 8/8 *fruM* males.

Uniquely among insects, mosquito ears contain an auditory efferent system that can modulate JO neuron properties^9^. The efferent system is composed of descending neurons from the CNS that innervate different regions of the JO (Figure 3B)^10^. The localization of efferent presynaptic terminals within JO is sexually dimorphic^33^. We therefore tested if FruM influences male-specific localization of presynaptic terminals. To visualize presynaptic terminals, we stained JO sections with presynaptic marker antibodies 3C11 and nc46^9^.

Following previous classification guidelines, we successfully identified all five types of presynaptic terminals in our JO sections^10^. We additionally observed presynaptic terminals located on the inner segments of JO dendritic cilia, which we classified as type VI presynaptic terminals. By comparing control males and females, we observed female-specific type III presynaptic terminals and male-specific type II, IV, V and VI terminals (Figures 3C-3E, S9A and S9B). Type I terminals were observed in both control groups (Figures 3C-3E, S9A and S9B).

We observed differences in *fruM* male JO presynaptic terminal localization compared to control males: 1) male-specific type V terminals appeared less abundant in *fruM* males, 2) male-specific type VI terminals were not detected in *fruM* males and 3) *fruM* males gained female-specific type III terminals (Figures 3C-3E, S9A and S9B; red arrows: III (somata), orange arrows: V (basal plate), purple arrows: VI (inner JO dendritic cilia)).

Sexual dimorphisms in efferent innervation of JO neurons are associated with efferent modulation of hearing function^33^. The male efferent system controls the onset and cessation of male ear-unique self-sustained oscillations (SOs), which are large, monofrequent flagella vibrations reflecting excessive amplification of female flight sounds hundreds to thousand times above baseline, a phenomenon associated with enhanced JO neuron motility^30,33^. The injection of tetanus toxin (TeNT) blocks efferent signaling by inhibiting the presynaptic release of neurotransmitters, triggering SO onset only in males^33^. Following TeNT injection, we found that only control males could show SOs, with *fruM* males not exhibiting any SOs (Figures 3F and S10A; Table S16).

We next injected 8-Br-cAMP, which can induce SOs by direct activation of JO neurons^12^. Again, we found that cAMP injection was able to induce SOs only in control males (Figure 3G; Tables S17 and S18), though injection increased the mechanical tuning frequency of all groups (Figure S10B; Tables S17-S19).

FruM promotes proper localization and formation of male-specific efferent presynaptic terminals within male JOs and inhibits formation of female terminals. FruM establishes and controls male ear SO machinery *via* direct modulation of JO neuron properties and/or indirect regulation of the JO-innervating auditory efferent system properties.

### FruM inhibits female-biased gene expression and promotes male-biased gene expression

To elucidate the transcriptional regulatory roles of FruM in shaping male-specific hearing function properties, we submitted pedicels for RNA-sequencing. Principal Component Analysis (PCA) showed that PC1 (42% of variance) was attributable to genotype (Figure 4A). The PCA plot indicates that despite lacking functional FruM protein (like control females), the overall gene expression pattern of *fruM* males displayed a hypermasculinized phenotype shifted away from both sexes.

**Figure 4:**
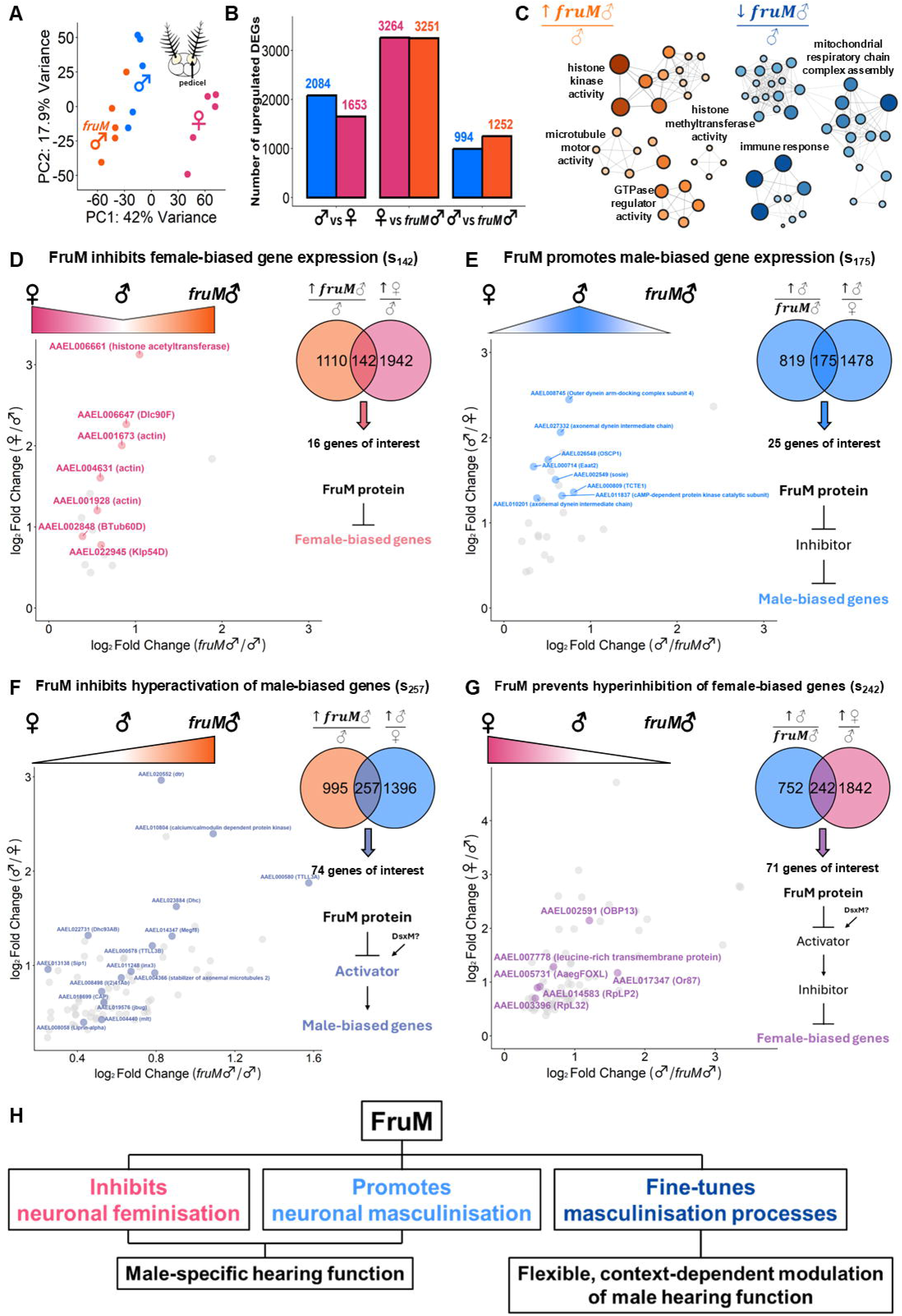
FruM-mediated transcriptional regulation. **(A)** Principal Component Analysis for control female, control male and *fruM* male pedicels. (Inset) Pedicel dissection diagram. Pedicels are colored in yellow. **(B)** Bar chart of differentially expressed genes across comparison groups based on DESeq2 analysis. **(C)** Network of GO enrichment terms identified as significantly enriched during analysis of genes significantly upregulated (left) or downregulated (right) in *fruM* males compared to control males. Size and color of the GO term nodes represent the number of genes identified under each GO term. Thickness of the edges represents the number of shared genes between two GO term nodes. See Figures S11A and S11B. **(D)** Log_2_ fold change expression of gene of interest in s_142_ that are upregulated in control females and *fruM* males compared to control males. Venn diagram shows number of overlapping genes between comparison groups. Highlighted are genes that show JO-specific and female-biased expression in published head snRNA-seq database^26^. See Figures S12A, S13A and S13B. **(E)** Log_2_ fold change expression of gene of interest in s_175_ that are upregulated in control males compared to control females and *fruM* males. Venn diagram shows number of overlapping genes between comparison groups. Highlighted are genes that show JO-specific and male-biased expression in published head snRNA-seq database^26^. See Figures S14A, S15A and S15B. **(F)** Log_2_ fold change expression of gene of interest in s_257_ that are upregulated in *fruM* males compared to control males and control males compared to control females. Venn diagram shows number of overlapping genes between comparison groups. Highlighted are genes that show JO-specific expression in published head snRNA-seq database^26^. See Figures S16A, S17A and S17B. **(G)** Log_2_ fold change expression of gene of interest in s_242_ that are upregulated in control males compared to *fruM* males and control females compared to control males. Venn diagram shows number of overlapping genes between comparison groups. Highlighted are genes with annotated function. See Figure S18A. **(H)** Schematic representation of FruM’s role in defining male hearing function.

Differential gene expression analysis between *fruM* and control males identified 1252 genes significantly upregulated in *fruM* males compared to control males and 994 downregulated genes (Figure 4B). Gene ontology (GO) enrichment analysis revealed that upregulated genes in *fruM* males had molecular functions related to histone modifications, microtubule motor activity and GTPase regulator activity (Figures 4C and S11A). Downregulated genes in *fruM* males were enriched with functions related to cellular respiration and mitochondrial activity, as well as immune response (Figures 4C and S11B).

To identify pedicel-specific genes potentially influenced by FruM, we: 1) identified significantly differentially expressed genes (padj<0.1 & log_2_FC>0 (upregulated) or log_2_FC<0 (downregulated)) (Tables S20-S23), 2) filtered for genes with higher expression in pedicels than heads in either sex using published head and pedicel RNA-seq databases^27^ and 3) examined expression patterns in putative JO neurons using published snRNA-seq database^26^.

Since both *fruM* males and control females do not produce functional FruM compared to control males, we started by searching for genes differentially expressed in the absence or presence of functional FruM. We first looked for genes significantly upregulated in *fruM* males and control females compared to control males and identified 142 genes (s_142_) (Figures 4D, S12A, S12B, S13A and S13B). In s_142_, we identified *histone acetyltransferase* (*HAT*) (AAEL006661) and *dynein light chain 90F* (*Dlc90F*) (AAEL006647) that showed strong female-biased expression in JO neurons (Figure S13B), suggesting their importance in shaping female-specific hearing machineries, and their expression could be under the inhibition of FruM in control male JOs^37,39–41^.

We further identified 175 genes significantly upregulated in control males compared to *fruM* males and control females (s_175_) (Figures 4E, S14A, S15A and S15B). In s_175_, we found eight genes with strong male-biased expression in JO neurons. This includes four genes (AAEL008745, AAEL027332, AAEL000809 and AAEL010201) that encode components of the ciliary motility apparatus that powers JO neuron motility, such as dynein intermediate chains and dynein docking and regulatory subunits^37^. Higher expression of s_175_ genes in control males could shape male-unique hearing function properties, such as male-specific SOs (hearing amplification) and enhanced mechanical/electrical sensitivity.

Given that expression of FruM in control males increased expression of s_175_ genes relative to other groups, we hypothesize that functional FruM could promote male-biased gene expression in control males by acting as a primary transcriptional inhibitor to inhibit expression of secondary transcriptional repressors that would otherwise inhibit male-biased gene expression (Figures 4D and 4E). The absence of functional FruM in *fruM* males and control females inhibits male-biased gene expression *via* elevated expression of intermediate transcriptional inhibitors (Figures 4D and 4E).

### Loss of fruM leads to an overall hypermasculinized gene expression landscape

In the PCA plot, we observed an overall hypermasculinized gene expression pattern in *fruM* males (Figure 4A), with hypermasculinized genes showing changes in expression from control females to control males to *fruM* males. We identified 257 genes that showed increases in expression following this pattern (s_257_) (Figures 4F, S16A-S16C, S17A and S17B). s_257_ contains genes that encode dynein heavy chains (AAEL022731 and AAEL023884) and dynein assembly factors (AAEL020552, AAEL004366 and AAEL008498), as well as genes that regulate microtubule dynamics (AAEL000580, AAEL000578 and AAEL004440). Given the importance of dynein and microtubule-related genes in powering JO neuron motility, we hypothesized that s_257_ could be essential in setting the genotype-dependent increase in ear tuning frequencies^35–37^.

The genotype-dependent increase in expression in s_257_ members could be shaped by antagonistic transcriptional activities of FruM and DsxM (Figure 4F)^20^. To establish a baseline increase in gene expression in both male groups compared to females, a masculinizing factor (such as DsxM) should transcriptionally promote expression of downstream transcriptional activators to increase male-biased gene expression (Figure 4F). However, to prevent excessive expression of male-biased genes in control males, FruM may act to inhibit expression of transcriptional activators to counteract DsxM’s activation, keeping expression levels of transcriptional activators in check (Figure 4F). The loss of functional FruM in *fruM* males could result in a lack of inhibition on downstream transcriptional activators; elevated expression levels of transcriptional activators in *fruM* males would subsequently result in elevated expression of male-biased genes in *fruM* males (Figure 4F). FruM thus prevents hyperactivation of male-biased genes in control males.

We identified 242 genes that showed decrease in expression from control females to control males to *fruM* males (s_242_) (Figures 4G and S18A). These genes were broadly related to ribosomal proteins and cellular respiration, potentially related to maintenance and turnover of cellular machineries in JO neurons^42–44^. Gene expression in s_242_ could be influenced by transcriptional inhibitors in s_257_ such that a negative correlation exists between expression of transcriptional inhibitors in s_257_ and female-biased genes in s_242_ (Figures 4F and 4G). Therefore, genotype-dependent increase in expression of transcriptional inhibitors in s_257_ from control females to control males to *fruM* males could result in a corresponding genotype-dependent decrease in female-biased gene expression in s_242_ (Figures 4F, 4G, S16B, S16C and S18A).

FruM thus fine-tunes masculinization processes by preventing both hyperactivation and hyperinhibition of male-biased and female-biased gene expression, respectively (Figure 4H)^45^.

### Motif enrichment analysis identified putative FruM-binding DNA motifs exhibiting in-silico interactions with FruM protein DNA binding domain

FruM proteins are putative transcription factors containing a BTB domain on their N-termini for protein-protein interactions and one of three C_2_H_2_ zinc-finger domains on their C termini for sequence-specific DNA binding^46–48^. Alternative splicing of *fru* transcripts at the 3’ end gives rise to different FruM protein isoforms (FruMA, FruMB or FruMC), depending on the zinc-finger domain incorporated^47^.

To identify putative FruM-binding genes that could validate the proposed FruM-dependent transcriptional networks, we focused on identifying FruM-binding motifs. We conducted motif enrichment analysis on 1kbp promoter sequences upstream of translational start sites (TSS) of genes in s_142_, s_175_, s_257_ and s_242_ (Figure 5A; Tables S24 and S25). The identified motifs showed minimal overlap between gene groups (Figure 5A). We focused on motifs identified in s_142_, as this gene group was predicted to be under the direct transcriptional inhibition of FruM. We conducted automated motif clustering based on sequence-similarity and found a cluster composed of three members whose consensus motif showed 7/10 base pair identity with the FruMB-binding motif in *Drosophila melanogaster* (*D. melanogaster*; Figures 5B and S19A)^20^. In control male pedicels, the *fru* exon encoding type B C_2_H_2_ zinc-finger domain (hereafter referred to as Zn-B domain) also showed the highest expression among all three zinc-finger coding exons. FruMB could thus be the predominant FruM isoform in male JOs (Figures 2B and S19B).

**Figure 5:**
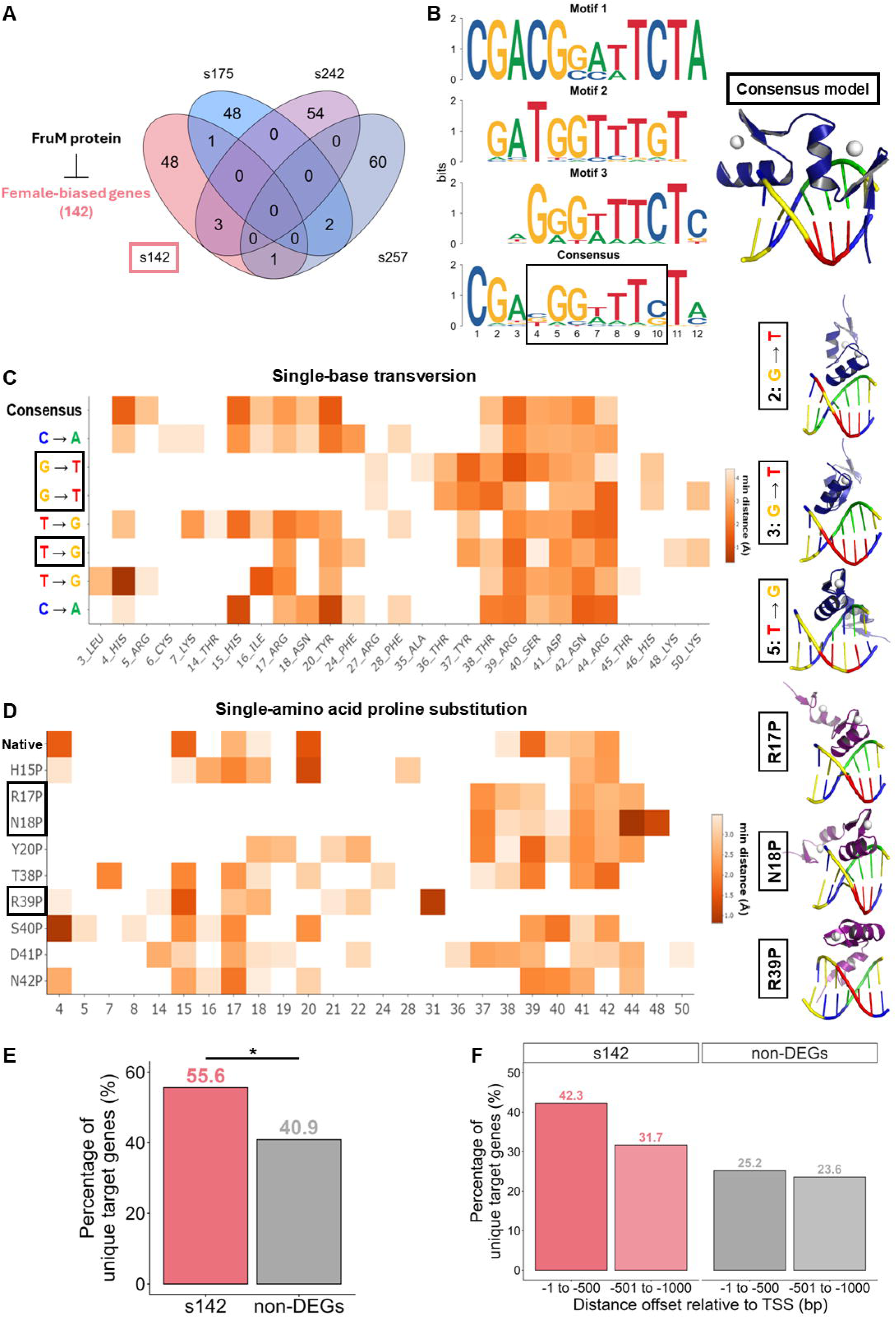
Identification of putative FruM-binding motifs. **(A)** Venn diagram showing overlapping motifs enriched in s_142_, s_175_, s_242_ and s_257_. **(B)** AlphaFold3 prediction of core consensus FruM-binding motif. Putative FruM-binding motif cluster identified based on automated motif clustering (left). Boxed nucleotide positions in the consensus motif are nucleotides that consistently showed interactions with Zn-B domain of FruM across all three members of the motif cluster. AlphaFold3 predicted protein-DNA complex between Zn-B of FruM and the seven core nucleotides of FruM consensus motif (right). See also Figures S19A-S19D. **(C)** Heatmap of the minimal distance (Å) between consensus or single-base mutated DNA motif and indicated amino acid residue of Zn-B. Only interactions with minimal distance < 4.5 Å were plotted (left). AlphaFold3 predicted protein-DNA complexes between Zn-B of FruM and the single base-mutated DNA motif for nucleotide position 2, 3 and 5 (right). **(D)** Heatmap of the minimal distance (Å) between the indicated amino acid positions of native Zn-B or single proline-substituted mutant protein and the consensus DNA motif. Only interactions with minimal distance < 4.5 Å were plotted (left). AlphaFold3 predicted protein-DNA complexes between consensus DNA motif and single proline-substituted mutant protein for amino acids R17, N18 and R39 (right). **(E)** Bar chart of percentage of unique target genes with at least one FruM consensus motif in the 1kbp promoter sequence upstream of translational start site for genes in s_142_ or non-differentially expressed genes between *fruM* male and control male pedicels (non-DEG group). Only identified motifs that showed conservation at positions 2, 3 and 5 with the consensus motif were included for analysis. *, p<0.05, Chi-squared test. **(F)** Bar chart of percentage of unique target genes with at least one FruM consensus motif within 500bp upstream of translational start site or 500-1000bp upstream of translational start site for genes in s_142_ or non-differentially expressed genes between *fruM* male and control male pedicels (non-DEG group). Only identified motifs that showed conservation at positions 2, 3 and 5 with the consensus motif were included for analysis.

We next used AlphaFold3 to predict interactions between the Zn-B of FruMB and candidate motif members^49,50^. We performed pairwise distance analysis and, across three models, consistently identified seven core nucleotide positions with < 4.5 Å interactions with amino acid residues of Zn-B, which we referred to as the FruM-binding consensus DNA motif (Figures 5B and S19C). To examine the specificity of these interactions, we performed single-base transversion (A↔C or T↔G) scanning across the seven positions and identified positions 2, 3 and 5 to be important positions mediating these interactions, with dissociation of interactions observed upon mutation (Figure 5C).

We found that among the 13 amino acids of Zn-B that showed interactions with the consensus DNA motif, 9 had interactions with minimal distance < 3.5 Å (Figure S19D). Four were part of the first alpha helix of Zn-B and five belonged to the second alpha helix, suggesting the importance of FruMB alpha helices in mediating sequence-specific recognition of consensus DNA motif (Figure S19D)^51,52^. We introduced single-amino acid proline substitutions for each of these amino acids to break the alpha helix structure. We identified Arg 17 and Asn 18 of the first alpha helix, and Arg 39 of the second alpha helix, to be respectively important for N and C termini interactions with the consensus DNA motif (Figure 5D).

Our results suggest that the C_2_H_2_ zinc-finger domain of FruMB could show sequence-specific binding with our putative FruM consensus DNA motif.

### Putative FruM consensus DNA motif enriched in promoter sequences of genes predicted to be under direct FruM transcriptional inhibition

To help validate our *in-silico* findings, we scanned for putative FruM-binding sites in the promoter region of genes in s_142_ and the non-differentially expressed gene (non-DEG) group (non-differentially expressed between *fruM* and control males) using the identified FruM-binding consensus DNA motif, with a strict criterion that positions 2, 3 and 5 of identified target sites should be identical to the consensus motif (Tables S24 and S26). We identified significantly more genes in s_142_ (55.6%) with at least one consensus motif in their promoter region compared to the non-DEG group (40.9%) (Chi-squared test; p < 0.05; Figure 5E). We also found a larger proportion of genes in s_142_ with consensus motifs located < 500 bp upstream of TSS compared to the non-DEG group (Figure 5F).

We next scanned for FruM consensus motifs in the promoter sequences of highlighted genes in s_142_ and identified female-biased genes including *Dlc90F* (AAEL006647) and *HAT* (AAEL006661) with at least one FruM-binding site, suggesting that FruM could bind directly to inhibit their transcription in control male JOs (Figures 4D, S12A and S19E; Table S24). We found two female-biased TFs *disco-R* (AAEL013374)^53^ and *mirr* (AAEL007502)^54^ in s_142_ that could be under the transcriptional inhibition of FruM (Figures 4D, S12B and S19F; Table S24). By inhibiting the expression of these TFs in control males, FruM could indirectly promote expression of male-biased genes in s_175_ (Figures 4D, 4E, S12B and S19F).

Lastly, we identified several transcriptional modifiers in s_257_ whose expression could be under direct inhibition of FruM (Figures S16B, S16C and S19G; Table S25). These transcriptional regulators include histone modifiers such as histone acetyltransferase *enok* (AAEL011106)^55^ and histone methyltransferase *trr* (AAEL019556)^56,57^, both of which increase gene expression by increasing chromatin accessibility, as well as two TFs, *Lim3* (AAEL007120)^58^ and *jumu* (AAEL018187)^59^ (Figures 4F, S16B, S16C and S19G; Table S25). These transcriptional activators could increase expression of downstream male-biased effector genes or further activate the expression of transcriptional inhibitors in s_257_ including *HDAC1* (AAEL004586)^45^, *lola* (AAEL009212)^60^, *XNP* (AAEL001141)^61^ and *Kr-h1* (AAEL002390)^62^, thereby suppressing expression of female-biased genes in s_242_ *via* transcriptional inhibition or induction of chromatin compaction (Figure S19G).

FruM exerts its transcriptional influence on downstream hearing gene expression *via* promoting or inhibiting the expression of intermediate transcriptional modifiers, with these processes tightly coordinated to define male hearing properties.

## Discussion

Strong sexual dimorphisms in mosquito reproductive behaviors have been documented for almost a century^3,4^. Males form mating swarms and find females *via* attraction to female flight sounds. Females do not display these behaviors, but instead seek hosts, blood-feed and lay eggs^3,4–7^. However, the molecular underpinnings of these behavioral dimorphisms remain unclear. Here, we show that the sex determination gene, *fru* is the master regulator of male reproductive behaviors. The loss of FruM completely abolishes male-specific aggregation and phonotaxis behaviors, resulting in mating failures of *fruM* males.

Male aggregation is a sex-specific rhythmic behavior under tight circadian regulation^5^. We observed that *fruM* males completely lack aggregation behavior, potentially due to disruption in sex-specific circadian rhythms (Figures 1B-1F). FruM may influence these circadian rhythms, possibly *via* co-expression or synaptic interactions with circadian clock neurons^63,64^. Support for the former is the hyperactivation of *cycle* expression in *fruM* males compared to control males in the absence of FruM (s_257_, Figures S16B and S16C). Future work could focus on examining if such interactions underlie rhythmic swarming behavior in male mosquitoes. Interestingly, we found that FruM does not regulate sexually dimorphic WBFs (Figure 1G), suggesting that other sexually dimorphic factors (such as *dsx*) could regulate sex-differences in baseline WBFs^65^.

Our work provides strong evidence that *fruM* male mating failures are due to a complete loss of phonotaxis behavior (Figures 1H-1I). Importantly, our results highlight that the loss of phonotaxis in *fruM* males could be attributed to primary defects in mechanical feedback amplification in hearing, sound detection and auditory information processing in mutant ears. We found that FruM is essential in setting the sensitivity and selectivity of male ears to female flight sounds. Our study also demonstrates that the mosquito peripheral auditory system represents a powerful model to study the actions of *fru* in an anatomically and functionally dimorphic sensory system. This opens up novel research avenue as most *fru* studies in *D. melanogaster* have focused on *fru*’s role in the central nervous system, and not the peripheral nervous system^48^.

Unlike *Ae. aegypti*, the ears of *D. melanogaster* are neither anatomically nor functionally dimorphic^30–32,35,37,66,67^. Here, we identified strong *fru* expression in JO neuron axons projecting to AMMC, suggesting FruM plays an indispensable role in male auditory processing relative to other sensory modalities. Furthermore, this intense *fru* expression stands in stark contrast to the sparse expression patterns observed in *D. melanogaster*^29^, indicating a far greater role of FruM in fundamentally defining the sexually dimorphic bases of hearing function, and thus the influence of male hearing on courtship outcomes, in mosquitoes. Given these species diverged ∼250 million years ago, it would be of interest to explore the evolutionary forces underlying such divergence in antennal ear dimorphic properties, and to understand how *fru* has acquired a new role to regulate sexually dimorphic neuronal differentiation in anatomically dimorphic mosquito ears^29^. Future work on multi-omics analysis of the antennal ears across mosquito and *Drosophila* species could help to address this knowledge gap.

We observed that localization of presynaptic terminals within the JO was altered in *fruM* males compared to control males, with mutant males displaying a lack/reduction of male-type and gain of female-type presynaptic terminals (Figure 3). Our work could not elucidate the underlying *fru*-dependent mechanisms that shape sexually dimorphic efferent neuronal arborization patterns within the JO, partially due to our lack of knowledge of the origins of these efferent neuronal populations, i.e. where these efferent neurons descend from. In *D. melanogaster*, *fru* shapes sexually dimorphic neuronal arborization patterns *via* axon guidance, tracking of axon-bound receptors along morphogen gradients of guidance cues or sex-specific programmed cell death during nervous system development^16,18,68–70^. We hypothesize that defects in these mechanisms could underlie the altered organization of auditory efferent terminals in *fruM* male JOs. The unique auditory efferent system of mosquitoes is a suitable model for studying these mechanisms, with future work needed to investigate at high resolution and across developmental stages the FruM-dependent mechanisms that are responsible for promoting localization and formation of male-specific efferent presynaptic terminals in male JOs.

Here, we elucidated key hearing genes and transcriptional modifiers that could be under the transcriptional influence of FruM (Figure 4). We attempted to further identify direct FruM-binding genes *via in-silico* protein-DNA interaction analyses (Figure 5). These promising targets represent the first step towards elucidating the transcriptional actions of FruM in mosquito ears. Future work could supplement our findings by conducting ATAC-seq or ChIP-Seq profiling to validate the identified protein-DNA interactions. Functional elucidation of FruM target genes could help to improve and refine the proposed FruM-dependent transcriptional regulatory networks. In one of our proposed models, DsxM and FruM share target genes, with the former acting as a transcriptional activator of gene expression while the latter inhibits. Future work could focus on elucidating if these two important male sex determination transcription factors act antagonistically to maintain appropriate levels of neuronal masculinization.

In summary, we propose FruM as the master regulator of male *Ae. aegypti* reproductive behaviors, including male aggregation and phonotaxis. FruM genetically defines the male-specific hearing system in mosquitoes, potentially the most important sensory system involved in the early stages of mosquito mating. *Via* sophisticated transcriptional networks, FruM modulates neuronal masculinization of male ears by striking careful balance between promoting male-biased and inhibiting female-biased gene expression programs, enabling male ears to be highly sensitive to female flight sounds. We propose *fru* as a new target for genetic-based control strategies aiming to exploit the sex determination pathway to control mosquitoes^71^.

## Resource availability

### Lead contact

Requests for further information and resources should be directed to, and will be fulfilled by, the lead contact, Matthew P Su.

### Materials availability

This study did not generate any new reagents. Requests for the *fruM* mutant lines should be addressed to Leslie Vosshall, with provision of the line pending scientific review and a completed material transfer agreement.

### Data and code availability

- Data are publicly available at the paper’s Mendeley data repository (available upon publication).
- All original code is available *via* Mendeley data (available upon publication).
- Any additional information required to reanalyze the reported data is available from the lead contact upon request.

## Acknowledgments

We would like to thank Mika Nomoto, Yasuomi Tada, Akiko Akama, and Mikako Yamaguchi (Center for Gene Research, Nagoya University) for assistance with RNAseq experiments at the Division for Medical Research Engineering. We thank Yixiao Zhang for technical support with mosquito rearing. We thank Ryoya Tanaka for his comments and suggestions and Nutnicha Kanjanapituk for the schematic diagram of a mosquito head. We thank Theo Maire for his advice regarding the BuzzWatch system. We finally thank Nipun Basrur and Leslie Vosshall for providing the *fruitless* mutant lines tested in this paper.

This study was funded by the following grants:

JST FOREST JPMJFR2147 (AK)

MEXT KAKENHI Grant-in-Aid for Transformative Research Areas (A) “Materia-Mind” JP24H02200 (AK)

Human Frontier Science Program Organization RGP0033/2021 (AK)

MEXT KAKENHI Grant-in-Aid for Research Activity Start-up JP22K15159 (MPS)

Nagoya University Tokai Pathways to Global Excellence 0121an0002 (MPS)

JSPS Invitational Fellowships for Research in Japan (Short-term) S22091 (DFE)

International Principal Investigator (PI) Invitation Program, Nagoya University, Japan (DFE) JSPS PhD 2524KJ1285 (YML)

## Author contributions

Conceptualization YML, DFE, MPS, AK

Data Curation YML, MPS, AK

Formal Analysis YML, AJT, MPS

Funding Acquisition YML, DFE, MPS, AK

Investigation YML, AJT, TTL, YYJX, WTC, MPS

Methodology YML, AJT, DFE, MPS

Project Administration YML, DFE, MPS, AK

Resources YML, DFE, MPS, AK

Software YML, MPS

Supervision MPS, AK

Validation YML, MPS

Visualization YML, MPS

Writing – Original Draft YML

Writing – Review and Editing YML, DFE, MPS, AK

## Declaration of interests

The authors declare that they have no competing interests.

## Materials and methods

### Animals used in this study

Unless otherwise noted, all mosquitoes tested were 5-7 days old. Both males and females were provided a diet of sugar water and tested females were not blood fed. Mosquito lines tested in this study were wild-type *Ae. aegypti* (Liverpool strain) and *Ae. aegypti fruM* mutant lines and background controls (Vosshall lab)^21^. All adults tested in this study were virgins. *fruM* males referred to in this study were heteroallelic null mutants generated from crossing *fruitless*^Δ*M*^ and *fruitless*^Δ*M-tdTomato*^ heterozygous parent lines. Male and female mosquitoes were used for all experiments described.

### Mosquito rearing

Mosquitoes were reared at 28°C and 60-70% relative humidity under 12 h:12 h Light:Dark (LD) cycles. Larvae were provided with fish food and pupae were sex separated. Adults were provided with 10% glucose water. Blood feeding was conducted *via* a membrane feeding system (Orinno Technology Pte Ltd., Singapore).

### Experiment entrainment paradigm

Adult mosquitoes (3-5 days old) were housed in cages or vials and entrained for two days under a 12:12 LD regime, except for the phonotaxis assay which adopted a 13:11 LD regime with 1 h of light ramping at the start and end of each light period. Experiments were carried out on entrained mosquitoes starting from the third day of entrainment. All experiments were conducted between Zeitgeber Time (ZT)11-ZT13, corresponding to the ‘dusk’ phase of entrainment. Mosquitoes had constant access to 10% glucose throughout. All experiments were either carried out in the entrainment incubator set to 22°C or performed in a temperature-controlled room at 22°C (±1.5°C).

### Locomotor activity monitor assay

Individual mosquitoes were aspirated into vials with glucose-soaked cotton wool provided on one end. Vials were then placed in LAM25 Locomotor Activity Monitors (Trikinetics). Following two days of entrainment in an incubator, the number of beam breaks per minute were counted for the next 72 hours. Analyses used the Rethomics package^81^.

Sample sizes: 40 control females; 49 control males; 58 *fruM* males.

### Video recording of flight behavior

Cages of 30 adult mosquitoes were entrained in an incubator for two days. Flight activity was recorded using a video camera (DMK33UP1300, ARGO; IC Capture Image Acquisition software ver 2.5) on the third day between ZT11.5-ZT12.

### BuzzWatch flight trajectory tracking assay

Groups of 30 adult mosquitoes were entrained in an incubator for two days. On the third day, flight activity between ZT11.5-ZT12 was recorded using a Raspberry Pi camera. Recordings were analyzed using the BuzzWatch analysis paradigm^24^ and heatmaps of mosquito position created based on analysis outputs. Trajectory points were filtered based on maximum flight speed, with points whose calculated speed was greater than 37.5 cm/s being dropped, and the remaining trajectory being split to create a new, separate trajectory. Maximum flight speed used for filtering was based on previous estimates of flight speed using the BuzzWatch system^24^.

To quantify the proportion of each trajectory spent close to the centre of the cage, the Euclidian distance from the centre to each point in the trajectory was calculated. The number of points at which the distance from the centre was less than 100 pixels was then divided by the total number of points included in the trajectory. Only those trajectories for which the proportion of time spent in the centre was greater than zero were included in the analysis. This meant that 250 (out of 915) control female trajectories, 392 (out of 481) control male trajectories and 232 (out of 673) *fruM* male trajectories were included in the quantification analysis.

Cage sample sizes: 3 control female cages; 3 control male cages; 3 *fruM* male cages. Trajectory sample sizes for quantification: 250 control female trajectories; 392 control male trajectories; 232 *fruM* male trajectories.

### Mosquito WBF and fly-by measurements

Cages of 30 adult mosquitoes were entrained in an incubator as described above. During ZT11-ZT12 on the third day, flight tones were recorded by putting a microphone (EK series, Knowles) into the cage. Recordings were made using a Picoscope 2408B and the Picoscope 6.14 software (Pico Technology) at 50 kHz sampling rate. The SimbaR package was used to analyze flight recordings ^5^ with only flight recordings longer than 150 ms included in the analysis. The median of all analyzed flight recordings was calculated. The number of fly-bys was calculated as the total number of analyzed flight recordings per cage.

Sample sizes: 11 control female cages, 11 control male cages and 11 *fruM* male cages.

### Group level phonotaxis assay

Male group phonotaxis assays followed published protocols^25,28^. Thirty adult male mosquitoes were aspirated into 30 × 30 × 30 cm insect cages (4S3030 Insect Rearing Cages, Bugdorm) and entrained in an incubator for two days. A speaker (FF225WK, Fostex) was placed next to the cage for playback of pure tones between 350 – 850 Hz (in 25 Hz bins). Pure tone files were generated in Audacity 3.0.3 with intensities calibrated to a Particle Velocity Level (PVL) of 1.58 × 10^−4^ ms^−1^, equivalent to 70dB Sound Pressure Level (SPL)^86^.

Males were tested for their phonotactic response to pure tones between ZT12.5-ZT13. Each tone lasted for one minute with one minute of silence provided between tones. The order of tone playback was pseudo-randomized between repeats using the sample function in R^87^. A video-camera (DMK33UP1300, ARGO) equipped with a lens (LM5JC1M, Kowa) was placed opposite the speaker to record phonotactic responses. The IC Capture Image Acquisition software (The Imaging Source, version 2.5) was used for all video recordings.

Quantification of phonotactic response was achieved by manual counting of the number of males landing on the netting directly next to the speaker during stimulation. The number of males located in front of the speaker immediately prior to tone onset was subtracted from the number which landed on the netting during tone playback. Counting was conducted without knowledge of tone frequency being played. Response counts were normalized within repeats.

Sample sizes: 2 control male cages; 2 *fru*^Δ*M*^/+ male cages; 5 *fruM* male cages.

### Video recording of individual phonotaxis

Four virgin control or *fruM* males were aspirated into a paper cup with glucose-soaked cotton inside. Mosquitoes were entrained for two days in an incubator and tested in a temperature-controlled room set to 22 ± 1.5 °C. Mosquito responses to 470 Hz pure tone were video recorded on the third day between ZT11.5-ZT12.

### Identification of target gene expression patterns using published snRNA-seq data

Adult *Ae. aegypti* head snRNA-seq data was downloaded from Goldman et al, 2025^26^ and expression patterns of target genes were plotted using the Seurat package in R^83^. Male and female cells were labelled based on the sex of the samples and putative JO neuron clusters were identified based on filtering for cells with *inactive* (AAEL020482) expression.

### *fru* splicing analysis

Differential splicing analysis of genes was performed using Dexseq^74^ in R. *Ae. aegypti* GFF file downloaded from VectorBase^85^ (68^th^ release) was used to create a flattened annotation. BAM files containing the aligned reads of *Ae. aegypti* male and female pedicels^27^ were mapped to the flattened annotation and exon usage counts for *fru* (AAEL024283) were obtained using the featureCounts function in Rsubread package^82^.

### JO immunofluorescence

Individual mosquitoes were sedated on ice before their heads were removed and fixed for 1 h in 4% paraformaldehyde (PFA) in Phosphate-buffered saline (PBS) with 0.25% Triton X-100 (PBT). Fixed heads were embedded in 10% agarose solution (Agarose S, 318-01195) and kept at 4°C for 10 mins to solidify the agarose blocks. Blocks were then fixed overnight in 4% PFA at 4°C.

Blocks were washed in 100% methanol for 10 min, transferred to PBS for 30 min and sectioned into 40 μm slices using a vibratome (Leica VT1200S). Sections were washed in 0.5% PBT three times at room temperature and blocked in 10% normal goat serum (NGS) (Vector Laboratories, Inc.)/0.5% PBT for 1 h. Sections were incubated at 4°C with primary antibodies in 10% NGS/0.5% PBT for three days.

Sections were washed three times with 0.5% PBT before a two-day incubation at 4°C with secondary antibodies in 10% NGS/0.5% PBT. After three washes with 0.5% PBT and one wash with PBS, sections were mounted on microscope slides and imaged with a laser-scanning confocal microscope (FV3000, Olympus) using 20× air objective (UPlanSApo, NA = 0.75) lens.

Primary antibodies used: anti-SYNORF1 3C11 monoclonal antibody (AB_528479, 1:30, Developmental Studies Hybridoma Bank (DSHB), University of Iowa); anti-SAP47 nc46 monoclonal antibody (AB_2617213: 1:30; DSHB); anti-DsRed polyclonal antibody (AB_10013483, 1:300, Takara Bio).

Secondary antibodies used: Alexa Fluor 488-conjugated anti-mouse IgG (A-11029, 1:300, ThermoFisher), Alexa Fluor 555-conjugated anti-rabbit IgG (A-21428, 1:300, ThermoFisher) and Cy3-conjugated goat anti-horseradish peroxidase (anti-HRP, RRID: AB_2338959, 1:150, Jackson Immuno Research).

### Brain immunofluorescence

Mosquito heads were fixed in 4% PFA in 0.25% PBT for 3 hours. Heads were washed twice with PBS before brains were dissected. Following dissection, brains were washed six times in 1% PBT at room temperature before permeabilization with 2% NGS in 4% PBT. After two days of incubation at 4°C, brains were washed six times with 1% PBT at room temperature then incubated in 1% PBT plus 2% NGS with primary antibodies for 2 days at 4°C. Brains were washed six times with 1% PBT at room temperature and incubated in 1% PBT plus 2% NGS with secondary antibodies for 2 days.

Finally, brains were washed six times with 1% PBT at room temperature and mounted in 80% glycerol in PBS. Samples were imaged with a laser-scanning confocal microscope (FV3000, Olympus) using a silicone-oil immersion 30× Plan-Apochromat objective lenses (UPlanSApo, NA = 1.05).

Primary antibodies used: anti-Brp (nc82) (AB_2314866, 1:30, DSHB) and anti-DsRed polyclonal antibody (AB_10013483, 1:300, Takara Bio).

Secondary antibodies used: Alexa Fluor 488-conjugated anti-mouse IgG (A-11029, 1:300, ThermoFisher), Alexa Fluor 555-conjugated anti-rabbit IgG (A-21428, 1:300, ThermoFisher).

### Mosquito preparation for laser Doppler vibrometry and electrophysiology

Mosquito preparation followed previous protocols^12,25^. Mosquitoes were sedated on ice and glued to a plastic rod using blue-light-curable glue (Norland Products Inc., 81). Only the mosquito’s right flagellum was free to move after gluing. Mounted mosquitoes were tested in a temperature-controlled room set to 22 ± 1.5 °C. The rod holding the mosquito was inserted into a micromanipulator (MM-3, Narishige Instruments, Japan) on a vibration isolation table. All vibrometry recordings were performed using a laser Doppler vibrometer (Vibroflex, Polytec) and the VibSoft software (Polytec).

For electrophysiology recordings, a sharpened tungsten reference electrode was inserted into the mosquito thorax to enable charging to −30 V relative to ground. Electrostatic actuators were positioned around the flagellum to enable provision of electrostatic stimulation. A tungsten recording electrode was inserted into the base of the pedicel to allow for antennal nerve recordings. The laser Doppler vibrometer was focused on the first flagellomere from the tip of the flagellum to enable simultaneous measurements of flagellar displacement. All electrophysiology recordings were collected using the Spike2 software (ver 10.08, CED), with stimuli created in Spike2.

### Laser Doppler vibrometry: Recordings

Stimulated recordings of the mosquito flagellum in the active state were made by measuring the mechanical vibrational response of the mosquito flagellum to white-noise (1–2000 Hz, Audacity 3.0.3) playback provided by a speaker (FF85WK, Fostex) placed 1.5 cm from the mosquito. The speaker was calibrated to a PVL of 1.58 × 10^−4^ ms^−1^, equivalent to 70dB SPL^86^.

For unstimulated recordings, prior to measuring the passive state free-fluctuations of mosquito flagellum, an active-state free-fluctuation recording was first obtained. Mosquitoes were then exposed to CO_2_ in a sealed chamber; depriving mosquitoes of oxygen leads to a (reversible) cessation of metabolic activity. An unstimulated recording of the sedated mosquito flagellar ear was immediately taken once CO_2_ delivery was stopped.

### Laser Doppler vibrometry: Data analysis

Fast Fourier transformation of raw time-domain data was conducted using the VibSoft software (Polytec) between 1 and 10000 Hz. Values below 125 and above 1000 Hz were excluded from further analyses due to noise. A forced damped oscillator function was fit to all transformed data using the lme4 package^77^ in R:

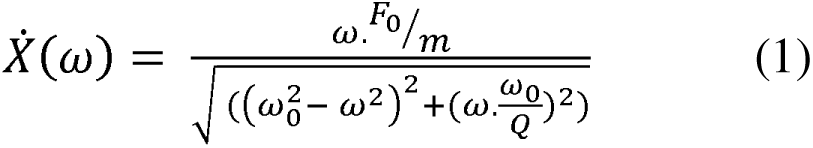

With F_0_ = external force strength, m = flagellar apparent mass, ω = angular frequency, ω_0_ = natural angular frequency and Q = quality factor = mω_0_/γ (γ = damping constant).

Fitting this function facilitated calculation of the natural angular frequency (ω_0_) for each recording, enabling calculation of the flagellar ear mechanical tuning frequency, f_0_, derived from f_0_= ω_0_/2π.

Sample sizes: 15 control females; 19 control males; 12 *fruM* males.

### Electrophysiology: Recordings

Prior to frequency sweep stimulation, a calibrated force-step stimulus was provided to each mosquito to calibrate the flagellum displacement to approximately ± 2.5μm. Stimulation paradigms followed published protocols^12,25^.

For frequency sweep experiments, sets of sweep stimuli, including one-second-long chirps of linearly increasing (forward, 1 to 1000 Hz) or decreasing (backward, 1000 to 1 Hz) frequency, were provided whilst the flagellar displacement (100 kHz sampling rate) and antennal nerve response (20 kHz sampling rate) were recorded in Spike2. Laser quality data (10 kHz sampling rate) was recorded to facilitate analyses.

For pure tone-based threshold experiments, 500 ms pure tone stimuli were provided at different levels of attenuation. Each series of pure tones per attenuation consisted of four sets of phasic and anti-phasic stimuli per frequency. Between each stimulus frequency, 500 ms of silence was provided, with one minute of silence between attenuations. Range of pure tones: 75 – 300 Hz for control females; 100 – 575 Hz for control and *fruM* males.

Mosquito flagella were exposed to a total of 40 sets of force step stimulation of seven different amplitudes. Each set of force step stimulation consisted of 200 ms positive force step and 200 ms negative force step with a 200 ms silence between directions. A 400 ms of silence was provided between two sequential sets of stimulation.

### Electrophysiology: Sweep analysis

For flagellar mechanical tuning analyses, laser quality data recorded as zero were excluded. Channel processes were applied to the laser channel data: a DC remove function (0.001s time constant); rectification; a smooth function (0.0005s time constant); a slope function (0.1s time constant). This enabled identification of when the channel slope was equal to zero, i.e. the time of maximum flagellar vibration. The corresponding stimulus frequency was taken as the peak ear mechanical tuning frequency.

For electrical tuning analyses of antennal nerve compound action potential responses, nerve responses to sequential phasic and anti-phasic stimuli were averaged to cancel out artifacts resulting from electrostatic actuation. The resulting nerve signal had the following processes applied: a DC remove function (0.01s time constant); rectification; a smooth function (0.0005s time constant); a slope function (0.1s time constant). Calculation of the time when the slope equaled zero enabled estimation of the corresponding stimulus frequency, referred to as the peak ear electrical tuning frequency.

For peak mechanical tuning frequency calculations, the average of consecutive phasic and anti-phasic sweeps of the same orientation was taken. The median of all these averages was taken as the peak mechanical tuning for that individual. For the peak electrical tuning frequency, the average between forward and backward sweeps was computed and the median was taken across averages.

Sample sizes: 19 control females; 17 control males; 13 *fruM* males.

### Electrophysiology: Pure tone analysis

Nerve responses to sequential phasic and anti-phasic stimuli were averaged to cancel artifacts in the nerve channel. A DC remove was applied to the resulting averaged nerve channel, followed by rectification. The Area Under the Curve (AUC) was then calculated for the 250 ms prior to stimulus onset and the last 250 ms of stimulus playback, for all stimuli. The threshold voltage for each stimulus frequency was defined as the voltage at which the AUC during stimulus playback was significantly greater than the AUC prior to playback, and for which all stimuli of greater voltage amplitude also elicited a significant nerve response. The minima threshold frequency was determined as the frequency that elicited significant antennal nerve responses at the threshold voltage. Nerve AUC response magnitudes were normalized to stimulus voltage for plotting. Maximum nerve phasic responses were determined as the peak nerve response estimated from loess fits of the nerve phasic response profile for each individual mosquito to pure tones provided at the largest stimulus voltage.

Sample sizes: 17 control females; 17 control males; 16 *fruM* males.

### Electrophysiology: Force-step stimulation analysis

Flagellar displacements in response to force-step stimulation were calculated by measuring the difference in flagellar displacements between the 5 ms average displacement at 100 ms after stimulus onset and the 5 ms average displacement immediately before stimulus onset. Nerve responses were calculated by subtracting 50 ms of average nerve baseline at 50 ms before stimulus onset from the nerve minima obtained within 2.5 ms after stimulus onset. Maximum nerve tonic responses were quantified as the largest nerve response recorded for each individual mosquito across different step magnitudes.

Sample sizes: 17 control females; 17 control males; 15 *fruM* males.

### JO width measurements

JO width measurements were calculated as the longest distance between the outer somata of both sides of the JO. Image stacks stained with HRP were used for measurements in ImageJ (version 1.53q)^88^. Only JOs in which the basal plate could be clearly identified were analyzed.

Sample sizes: 14 control females; 14 control males; 13 *fruM* males.

### cAMP and TeNT injection experiments

Mosquitoes were mounted as described above for vibrometry experiments. A free fluctuation recording was taken using the VibSoft software (Polytec) to act as a baseline prior to injection. Following this, compounds were injected into the mosquito and further recordings were made.

For cAMP experiments, either 3mM 8-Bromoadenosine 3,5-cyclic monophosphate sodium salt (8-Br-cAMP; 76939-46-3, Sigma), or Ringer’s control^12^ solution was injected into the base of the ear whose flagellar vibrations were not recorded from. Recordings were made 5, 10, 15 and 20 mins following injection.

For TeNT injection, 2μM TeNT (T3194, Sigma) or Ringer’s solution was injected into the mosquito thorax. Recordings were made every 5 minutes for 60 minutes following injection.

Recordings were analyzed as described above to extract the peak mechanical tuning frequency, enabling calculation of frequency change (ΔFrequency) at each recording timepoint compared to baseline. The largest ΔFrequency within an individual recording was taken as the maximum ΔFrequency.

Fluctuation power was calculated by first converting data to the displacement squared domain, then estimated using the AUC function in R. By dividing the AUC for each post-injection timepoint by the pre-injection baseline, the maximum AUC ratio following injection was calculated.

Sample sizes for 8-Br-cAMP / Ringer injections: 10/7 control females; 8/8 control males; 8/8 *fruM* males.

Sample sizes for TeNT / Ringer injections: 3/3 control females; 3/3 control males; 4/3 *fruM* males.

### Transcriptomics sample preparation

5-6 days old mosquitoes were flash frozen in liquid nitrogen at ZT12. Pedicels were dissected on ice in RNAiso Plus (9109, Takara Bio Inc.). Tissues were homogenized using a handheld homogenizer (BT LabSystems). RNAiso Plus was added to the homogenate to make up to a final volume of 1 mL and incubated at room temperature for 15 minutes prior to being added with 200 µL of chloroform (Kanto Chemical Co., Inc.). Samples were well-mixed followed by a 15-minute incubation at room temperature. Samples were centrifuged at 12,000g for 15 minutes at 4°C. After centrifugation, supernatant was transferred to a fresh tube followed by the addition of 0.5 mL of ice-cold 2-propanol (Sigma) to each sample. Samples were mixed well before being kept at −20°C for 30 minutes.

Samples were thawed and centrifuged at 12,000 × g for 15 minutes at 4°C, supernatant was discarded, and the RNA pellets were washed with 1 mL of 75% ethanol (Sigma) solution twice. RNA pellets were dried and dissolved in DEPC water (Nacalai Tesque). RNA quality was assessed using Nanodrop (ThermoFisher Scientific). Five biological repeats for each group were submitted to the Center for Gene Research at Nagoya University for library preparation.

### Transcriptomics: Library preparation, read alignment and differential expression analysis

*Ae. aegypti* L5 genome fasta file was downloaded from VectorBase (68^th^ release) for read alignment. Single-end reads were aligned to the genome using the align function in Rsubread in R. By using the *Ae. aegypti* GFF file downloaded from VectorBase (68^th^ release), gene counts of aligned reads were obtained using the featureCounts function provided in Rsubread package.

Gene annotation was conducted by retrieving *Ae. aegypti* gene relevant information from the EnzemblMetazoa using the biomaRt package^72^ in R. Differential expression analysis was carried out using DESeq2 package^73^ in R by setting a fold-change threshold of 1 and FDR cutoff <0.1. DESeq2 normalised counts were exported and used as RNA expression counts in heatmaps presented.

To further restrict the differentially expressed gene lists to pedicel-relevant genes, we applied a filter criterion for all analyzed gene lists to only include genes that showed higher summed read counts in the pedicel than in the head of either sex, by using a previously published pedicel and head transcriptomic datasets of male and female *Ae. aegypti* mosquitoes^27^.

### Aedes aegypti and Drosophila melanogaster gene ortholog conversion

Gene orthologs between *Ae. aegypti* and *D. melanogaster* were downloaded from OMA Browser^78^ and EnsemblMetazoa^89^ (downloaded on 7^th^ January 2025). EnzemblMetazoa ortholog gene list was fetched *via* the biomaRt package in R^72^. Duplicates from the two downloaded gene lists were removed. The ortholog gene list file was used to improve *Ae. aegypti* gene annotation.

### Improve *Ae. aegypti* GAF file for Gene Ontology enrichment analysis

Complete proteome of *Ae. aegypti* was downloaded from NCBI (downloaded on 7^th^ January 2025) and subjected to GO annotations using PANNZER2^79^. PANNZER2 results were combined with GAF file downloaded from VectorBase followed by removing duplicates.

### Gene Ontology enrichment analysis

The improved GAF file was used to run GO enrichment analysis using g:Profiler^75^. Custom GMT file used in this study can be found using the following g:Profiler token: gp E7yL_EL9T_8t8. The SCS threshold algorithm was selected for significance testing with a significance threshold of 0.05 applied.

### Motif enrichment analysis

Motif enrichment analysis on the 1kbp promoter sequences upstream of translational start site of target genes was conducted using findMotifs.pl (-fasta) from HOMER^76^. Background fasta files used to run the enrichment analysis included the 1kbp promoter sequences of all genes in the genome that were not part of the target gene groups. Promoter sequences were retrieved using the biomaRt::getSequence^72^ function in R. Motifs of interest were further scanned in the promoter sequences of other gene groups to determine their motif occurrences and positions using the findMotifs.pl (-find). Automated motif clustering was performed using universalmotif package^84^ in R.

### AlphaFold3 prediction of protein-DNA interactions

The C_2_H_2_ zinc-finger B protein domain (C_2_H_2_-Zn-B) of FruM was used as the protein query to predict interactions with target DNA motifs using AlphaFold3^49,50^. Two Zn^2+^ ions were also included to model the interactions. Predicted protein-DNA models were downloaded from AlphaFold3 and analyzed in PyMol^80^. The pairwise distance between interacting amino acid residues and motif base pair residues was extracted using the pairwisedistances.py script available on PyMolWiki. Minimal pairwise distance threshold of <4.5 Å was determined based on prior studies^90–92^. Single base pair mutagenesis was conducted *via* base pair transversion (C↔A or T↔G). Amino acid substitutions at alpha helixes were done by replacing target amino acid with proline to introduce kink in the alpha helix structure.

### Quantification and statistical analysis

P < 0.05 (prior to correction) was set as the significance level for all statistical testing. Two sided statistical tests were used throughout.

Shapiro-wilk tests of normality were first used to test for normality for all flight assay, functional assay and JO width data. Flight assay and JO width data were found to be normally distributed, so pairwise t-tests with BH corrections were used for statistical testing. Functional datasets including sweep data, as well as white noise stimulus and sedation vibrometry recordings, were found to be normally distributed; as such, pairwise t-tests with BH corrections were also used.

Functional datasets including threshold experimental data were found to be non-normally distributed, as well as datasets including proportions of flight trajectories within 100 pixels of the cage centre from the BuzzWatch analysis and vibrometry datasets for experiments focused on injection of ringer, cAMP and TeNT. Pairwise Wilcoxon tests with BH corrections were thus used.

---

**Table 1.** Parameters associated with flight and vibrometry for all groups tested. All values provided as means ± standard deviations. Numbers in brackets refer to sample sizes for each experiment.

| Group | Fly-by number | Wing Beat Frequency (Hz) | Peak mechanical tuning – WN (Hz) | Peak mechanical tuning – Sedated (Hz) |
| --- | --- | --- | --- | --- |
| Control female | 26.1 ± 12.1<br>(n = 11) | 467.8 ± 11.5<br>(n = 11) | 245.18 ± 14.63<br>(n = 15) | 234.37 ± 22.25<br>(n = 15) |
| Control male | 85.2 ± 23.6<br>(n = 11) | 751.5 ± 14.4<br>(n = 11) | 397.92 ± 20.38<br>(n = 19) | 324.40 ± 27.67<br>(n = 19) |
| *fruM* male | 21.1 ± 6.6<br>(n = 11) | 754.9 ± 25.0<br>(n = 11) | 488.50 ± 61.40<br>(n = 12) | 330.53 ± 29.79<br>(n = 12) |

**Table 2.**
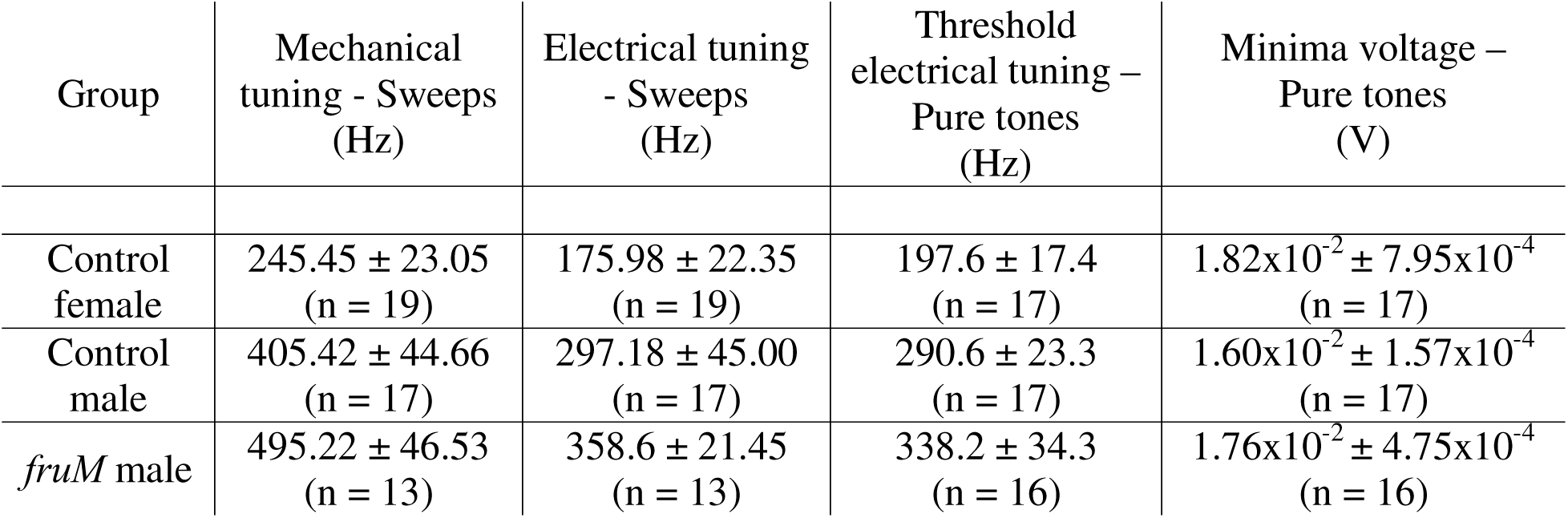
Parameters associated with electrophysiology for all groups tested. Sweep data provided as means ± standard deviations, pure tone data provided as median values ± standard errors. Numbers in brackets refer to sample sizes for each experiment.

## Notes

### Competing Interest Statement

The authors have declared no competing interest.

