## Supplemental Figures and Tables for "The sex determination gene *fruitless* is essential for male *Aedes aegypti* mosquito swarming and attraction to female flight sounds"

**Supplementary Figure 1: Loss of FruM alters male locomotor activity**

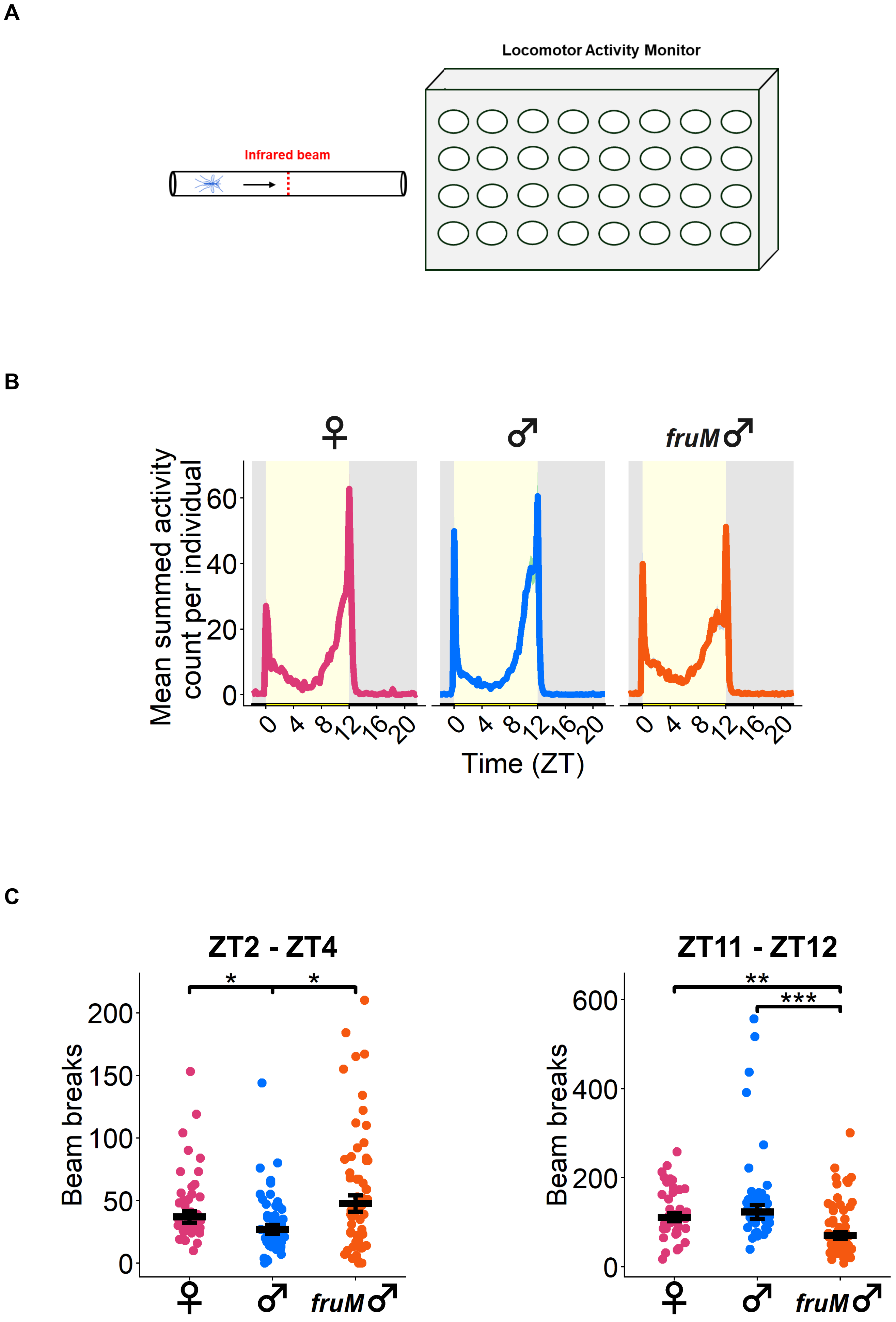

**Supplementary Figure 2: Heatmaps of mosquito flight activity at dusk**

**A**

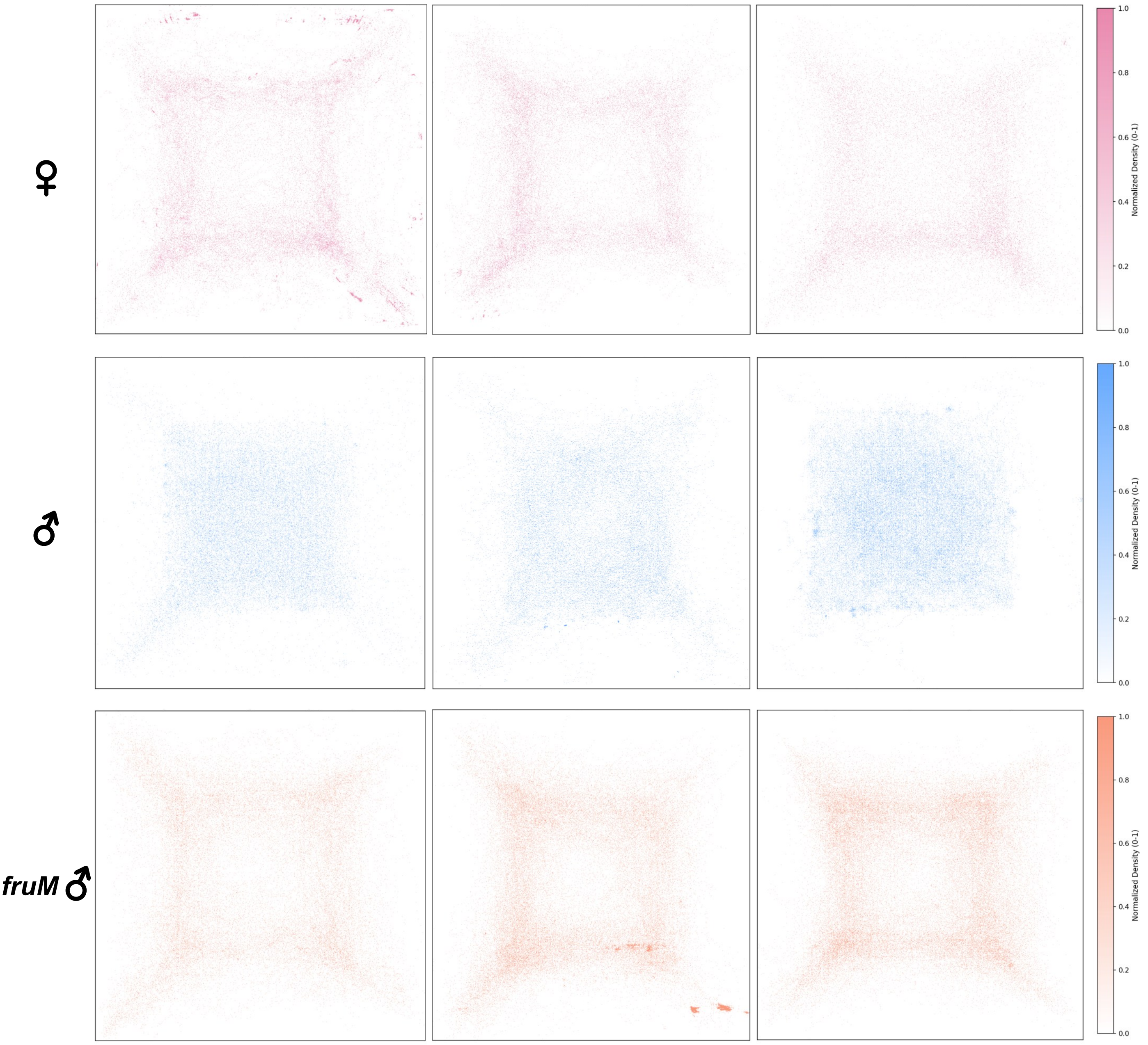

### Supplementary Figure 3: Control female flight trajectories from BuzzWatch recordings

**A**

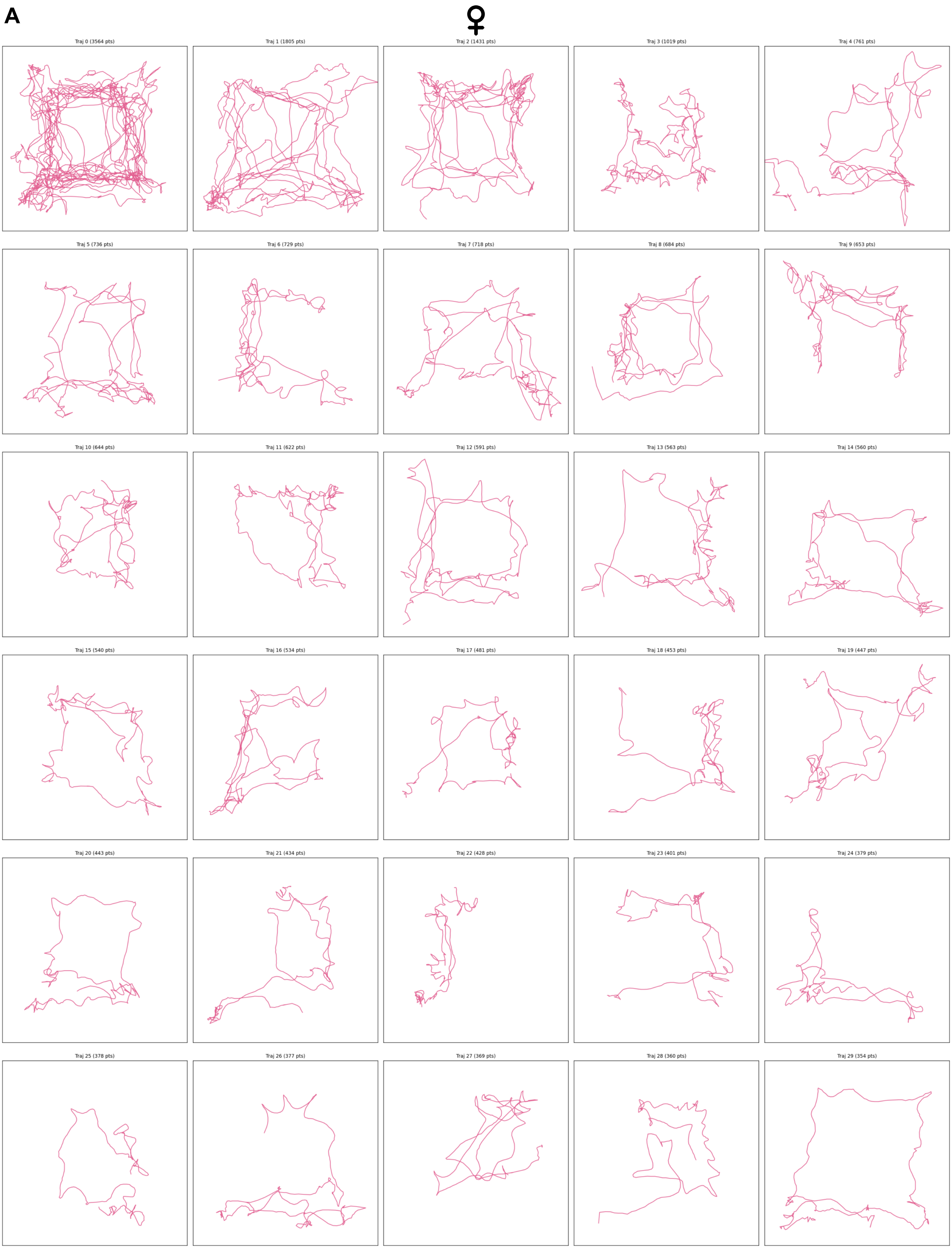

### Supplementary Figure 4: Control male flight trajectories from BuzzWatch recordings

**A**

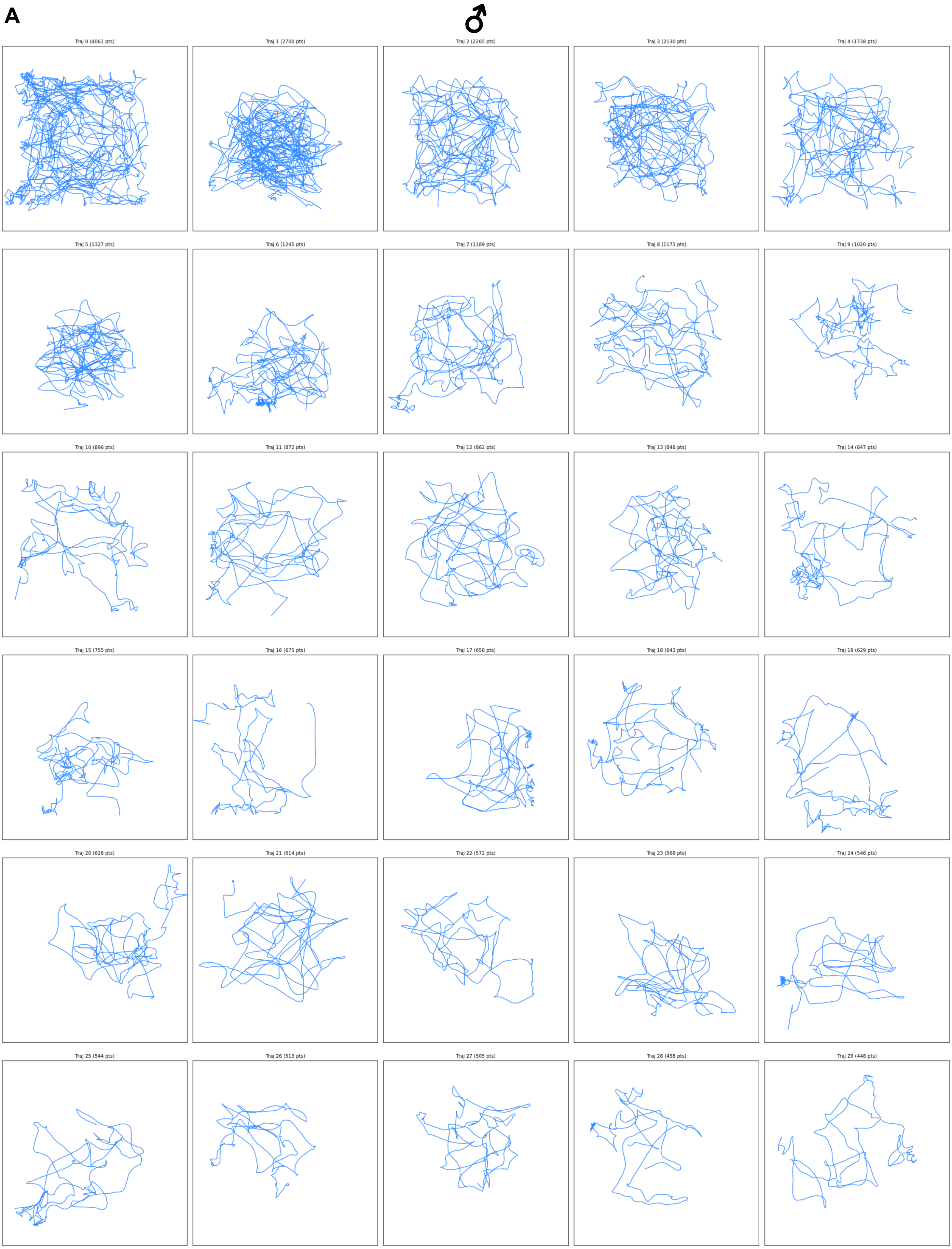

### Supplementary Figure 5: *fruM* male flight trajectories from BuzzWatch recordings

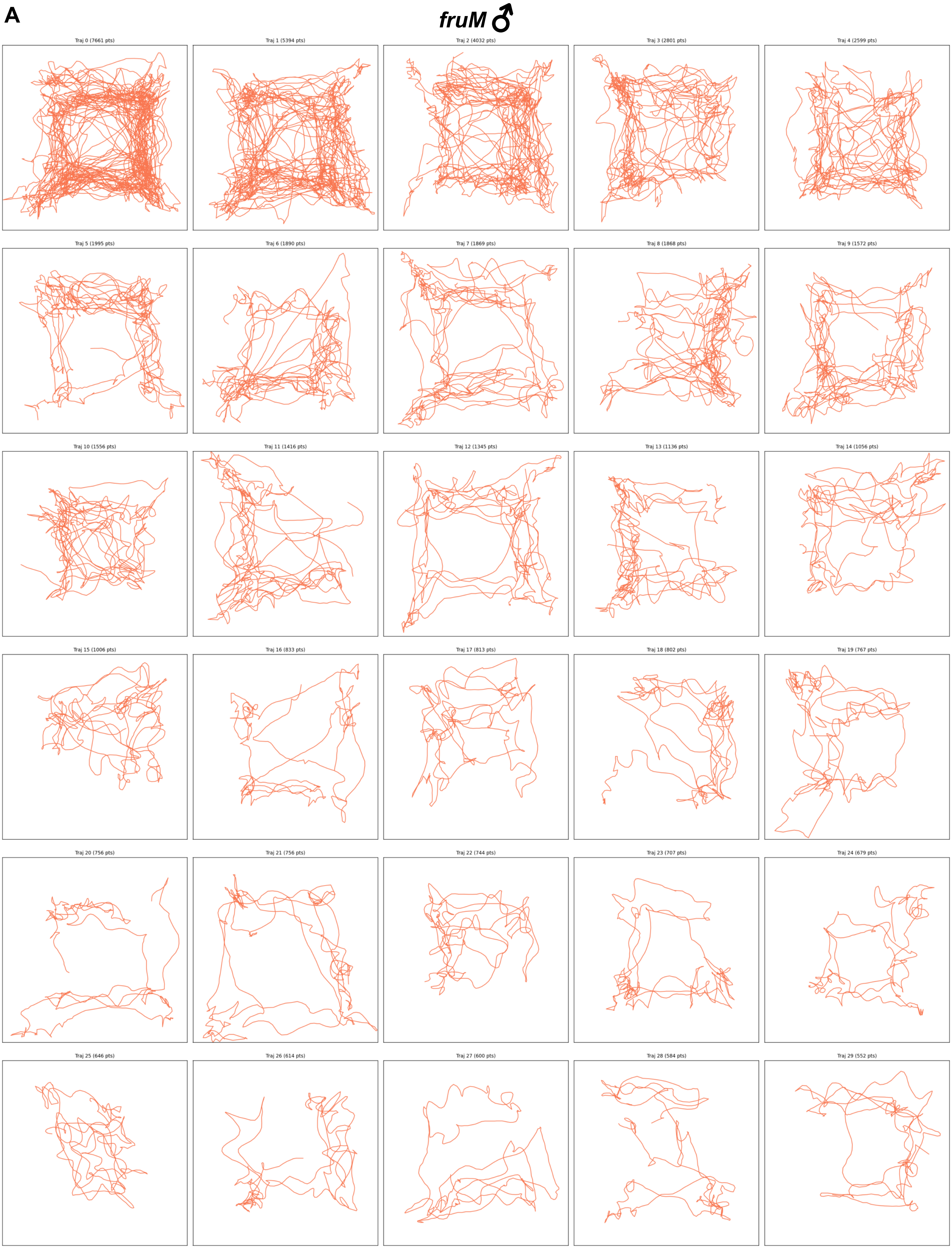

**Supplementary Figure 6: Loss of FruM abolishes male phonotaxis**

**A**

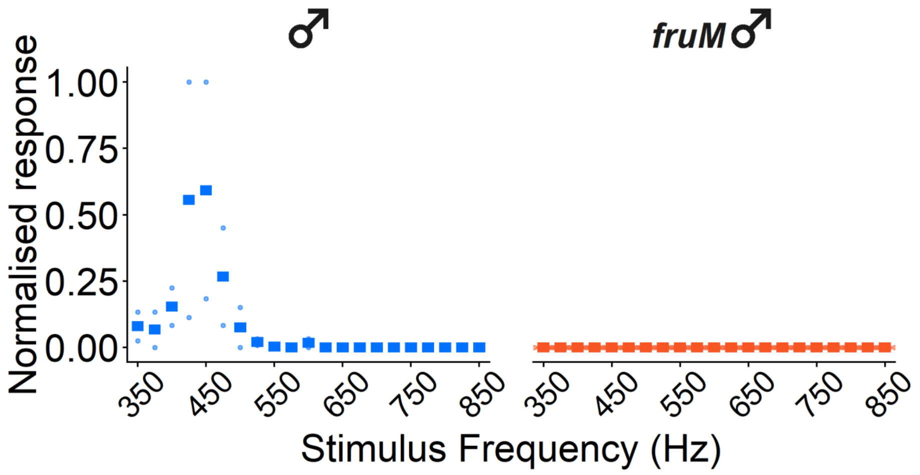

**B**

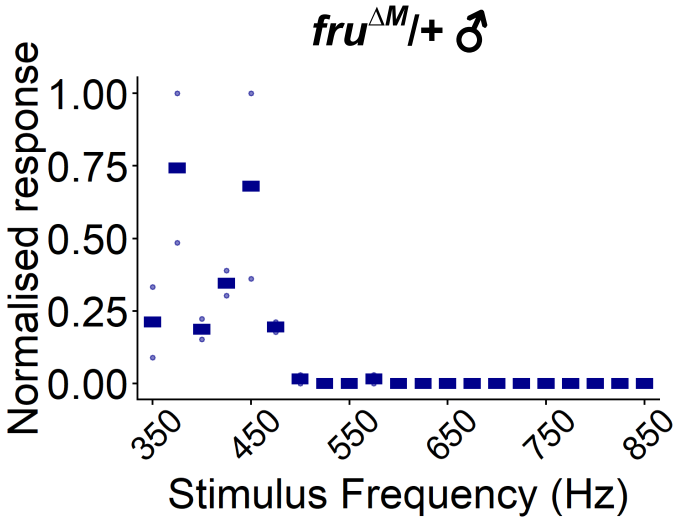

Supplementary Figure 7: *fru* expression in male JO and AMMC

A

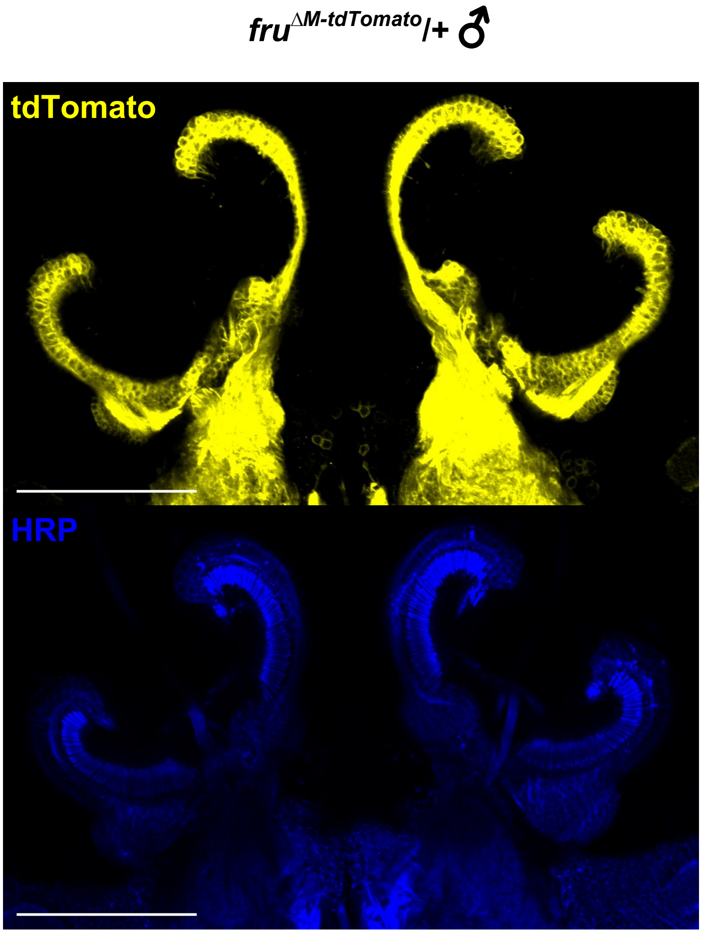

B

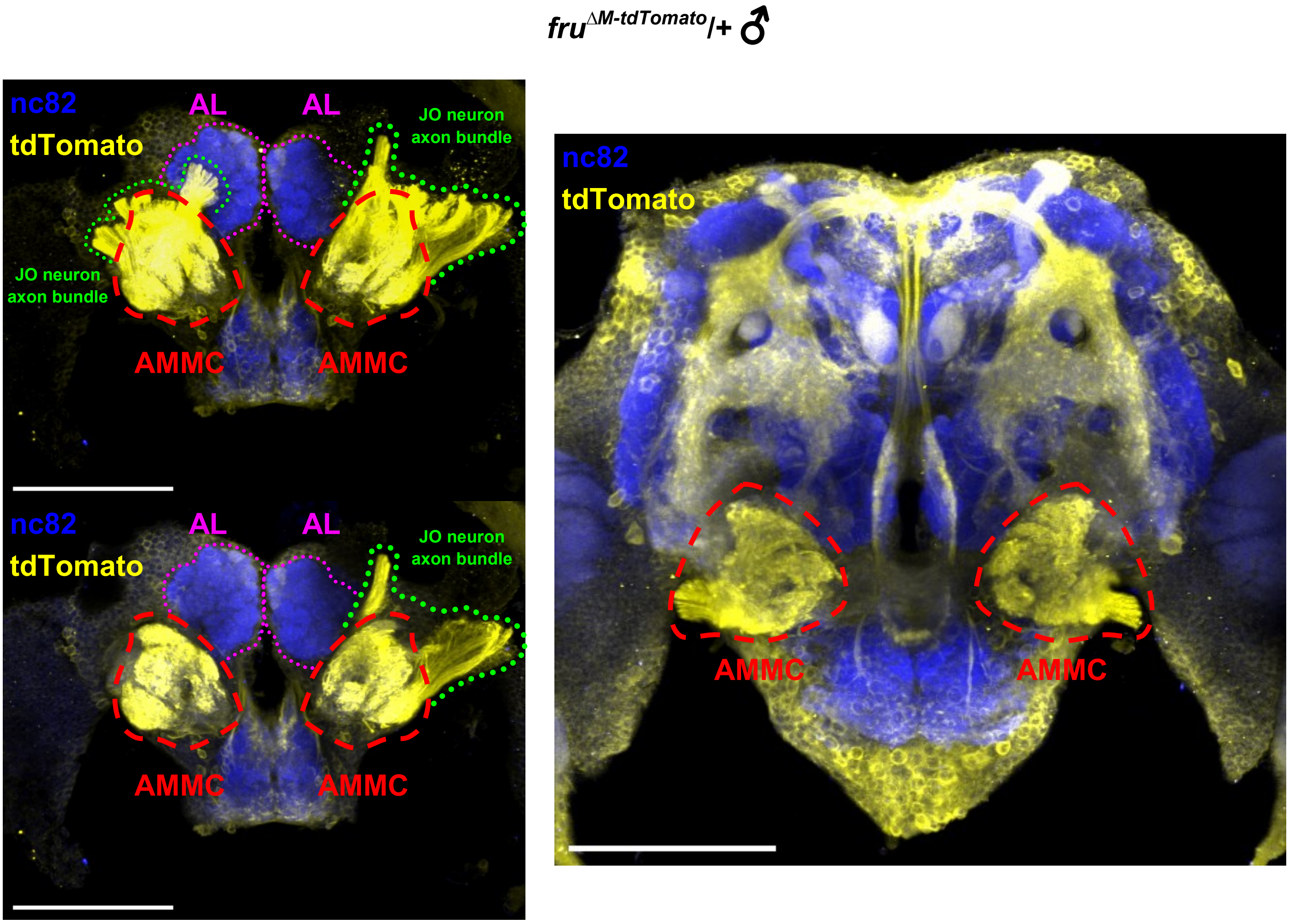

**Supplementary Figure 8: Loss of FruM alters male hearing function**

**A**

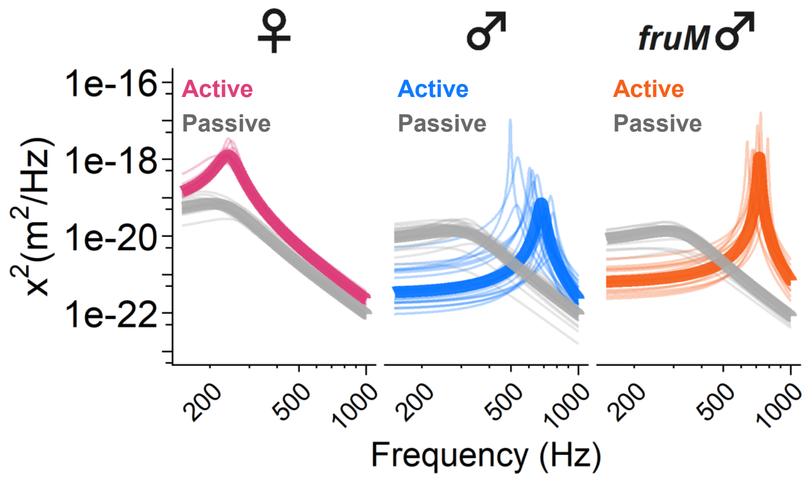

**B**

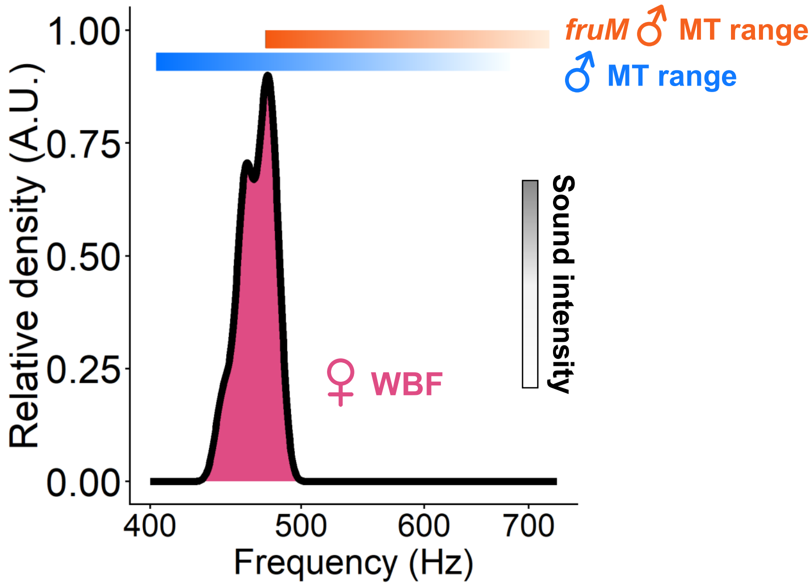

Supplementary Figure 9: Loss of FruM alters localization of presynaptic terminals in male JOs

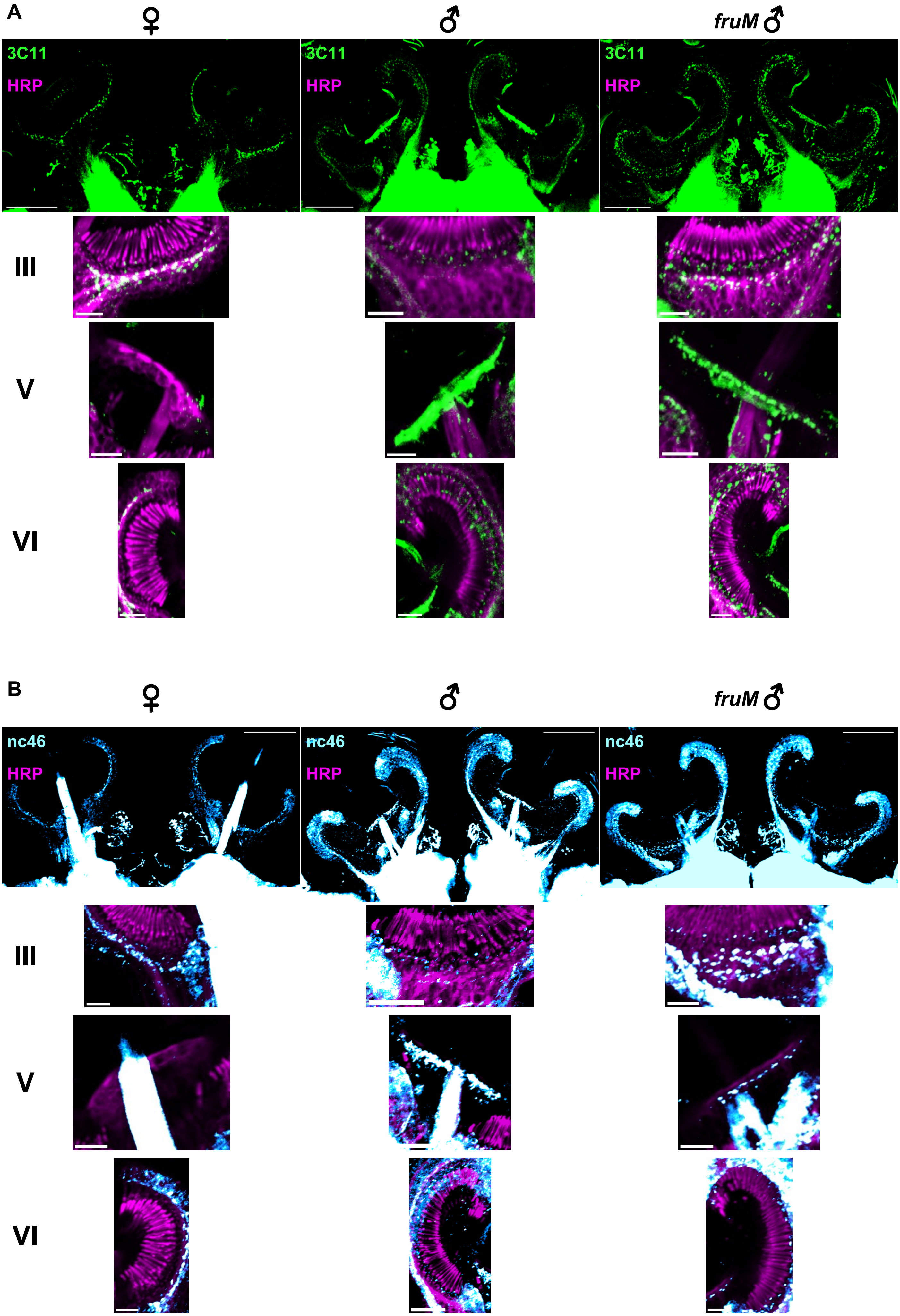

**Supplementary Figure 10: Changes in mosquito hearing function following compound injection**

**A**

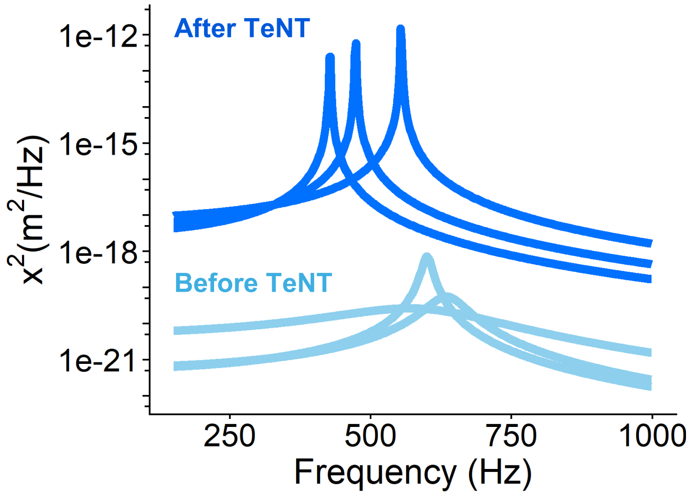

**B**

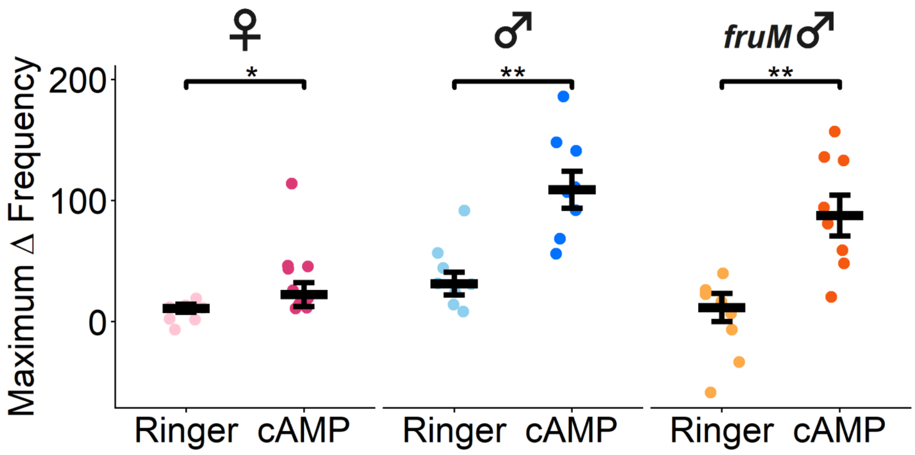

**Supplementary Figure 11: Gene ontology enrichment analysis**

**A**

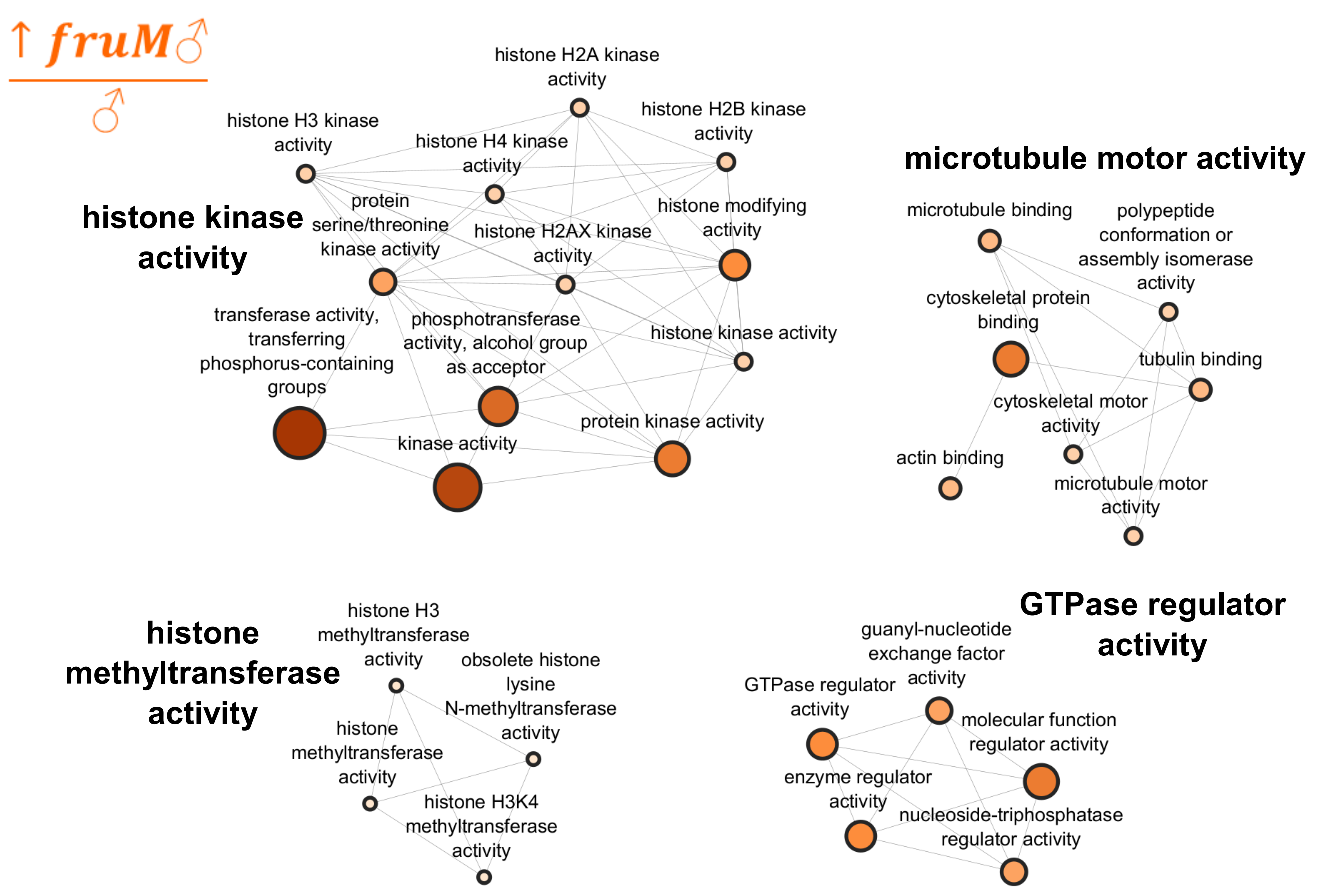

**B**

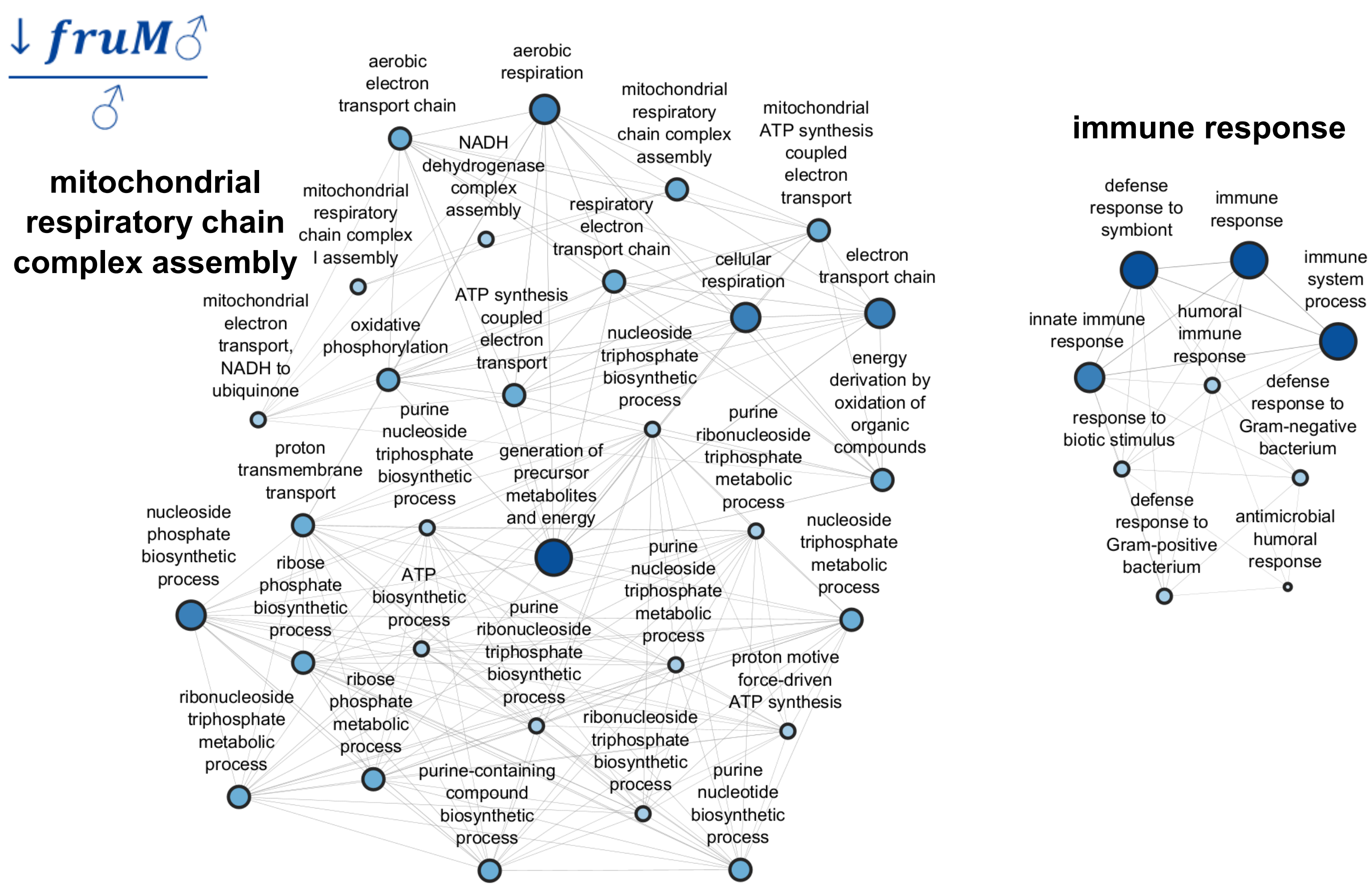

**Supplementary Figure 12: Expression levels of genes in $s_{142}$**

**A**

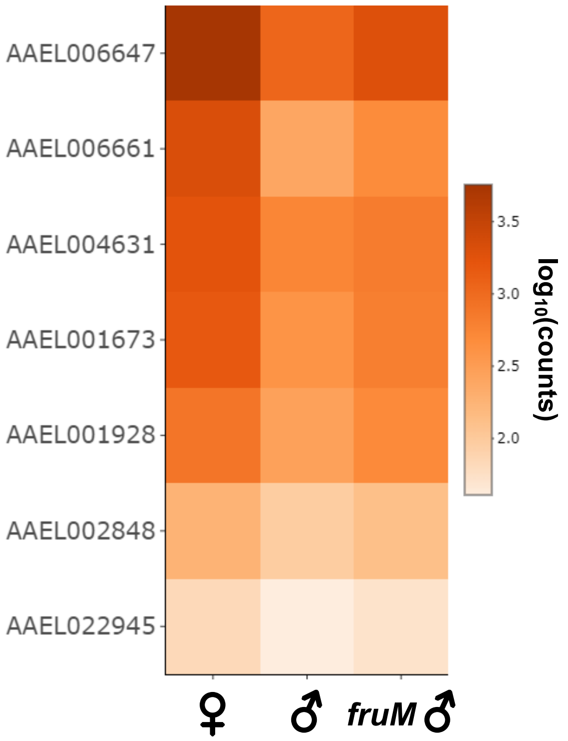

**B**

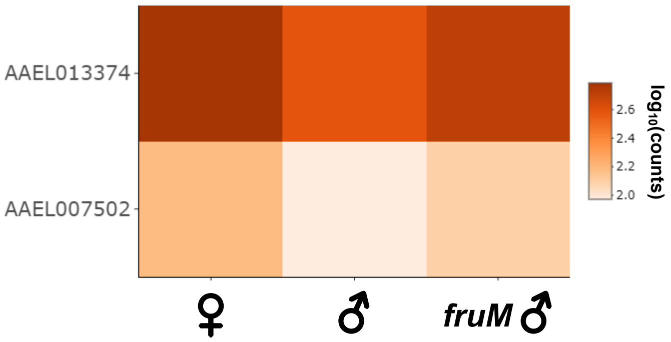

**Supplementary Figure 13: Expression of $s_{142}$ highlighted genes in Goldman et al, 2025 snRNA-seq database**

**A**

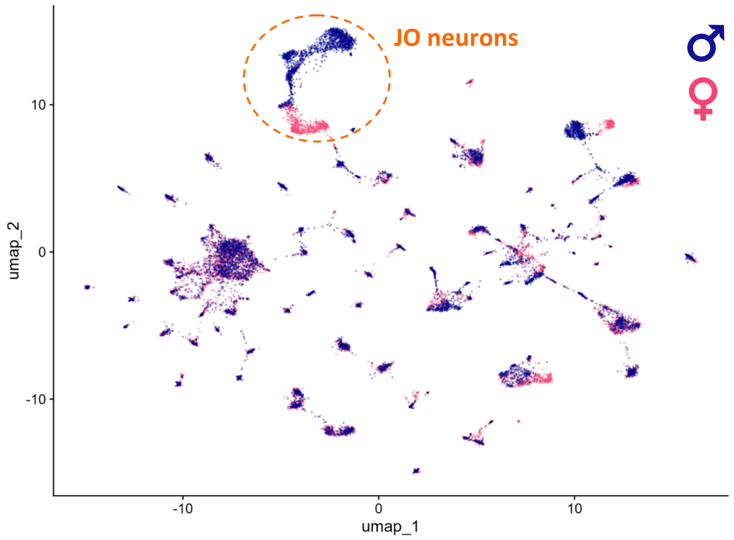

**B**

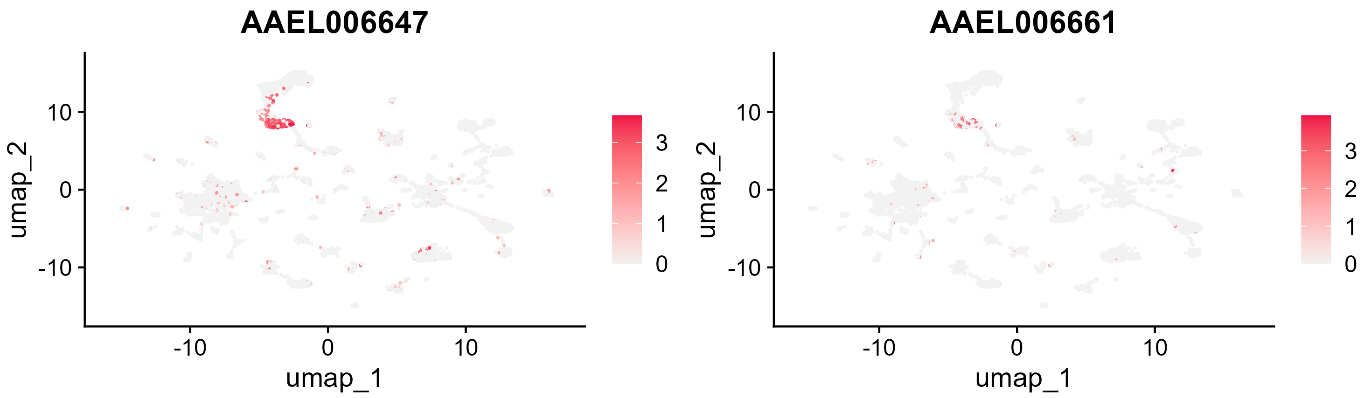

**Supplementary Figure 14: Expression levels of genes in $s_{175}$**

**A**

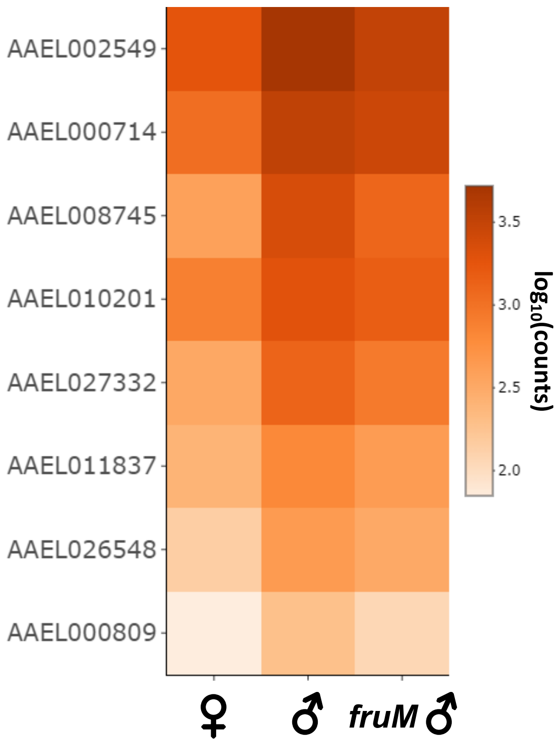

**Supplementary Figure 15: Expression of $s_{175}$ highlighted genes in Goldman et al, 2025 snRNA-seq database**

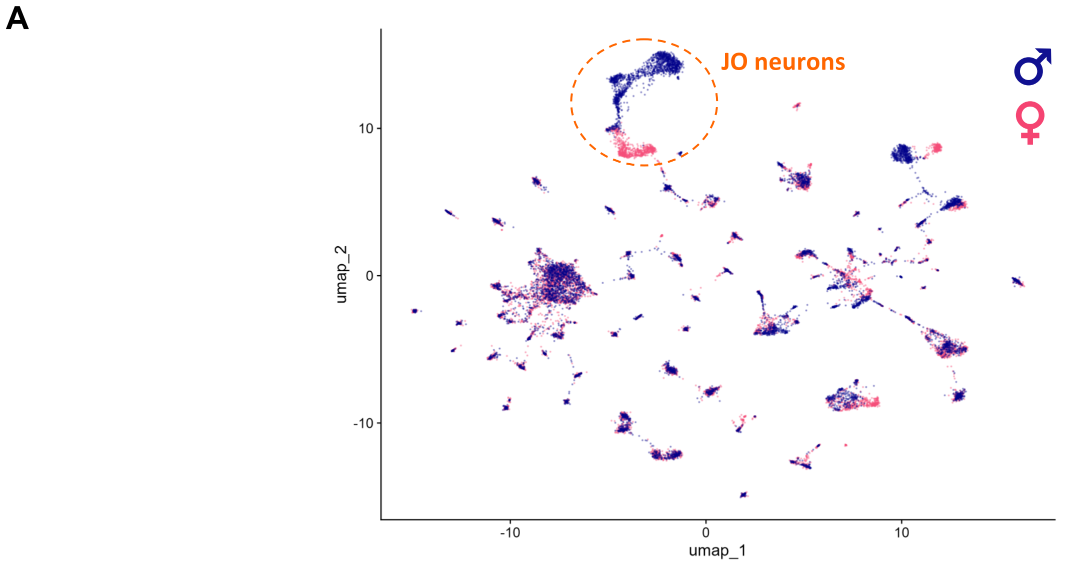

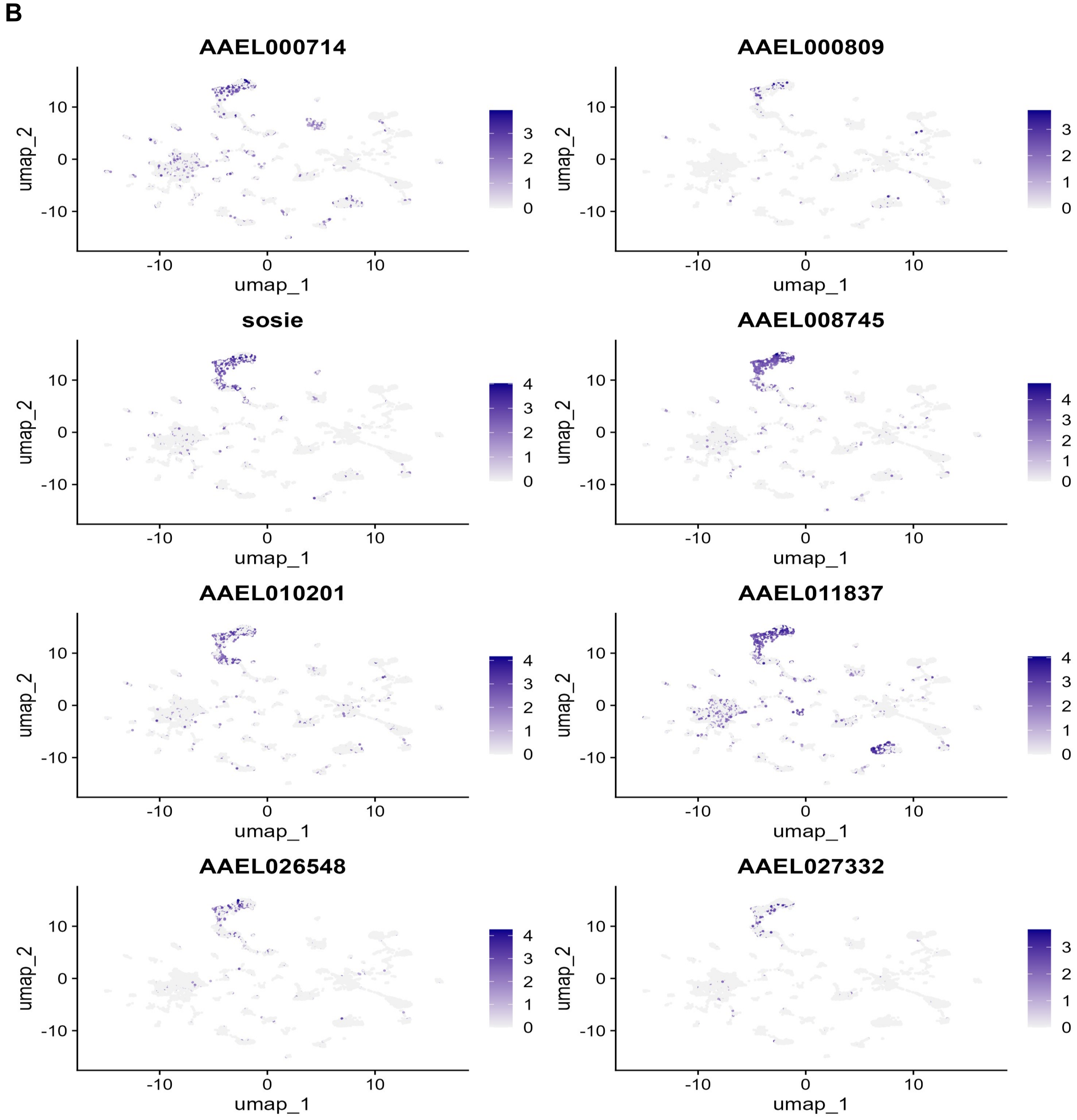

**Supplementary Figure 16: Expression levels of genes in $s_{257}$**

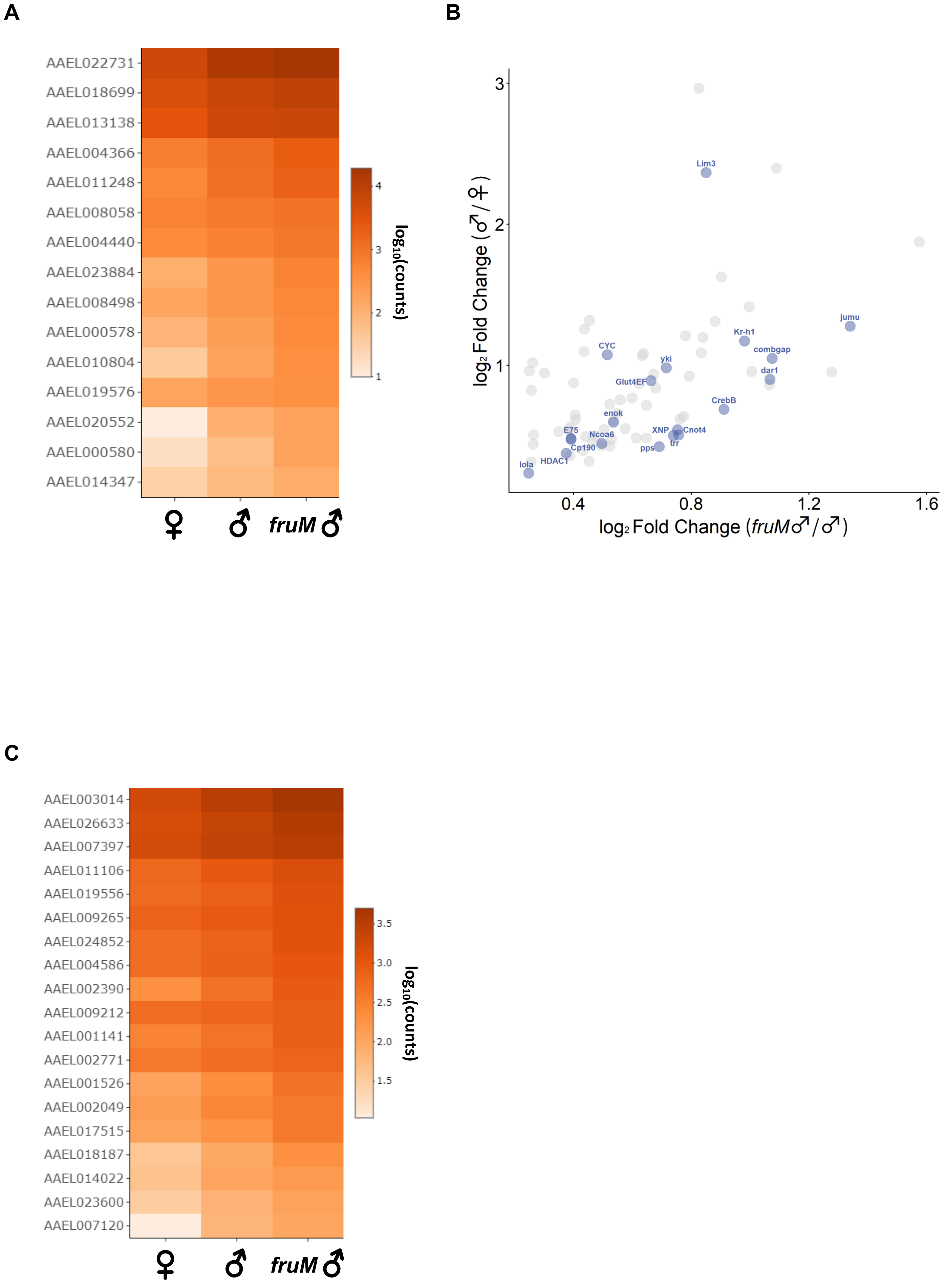

**Supplementary Figure 17: Expression of $s_{257}$ highlighted genes in Goldman et al, 2025 snRNA-seq database**

**A**

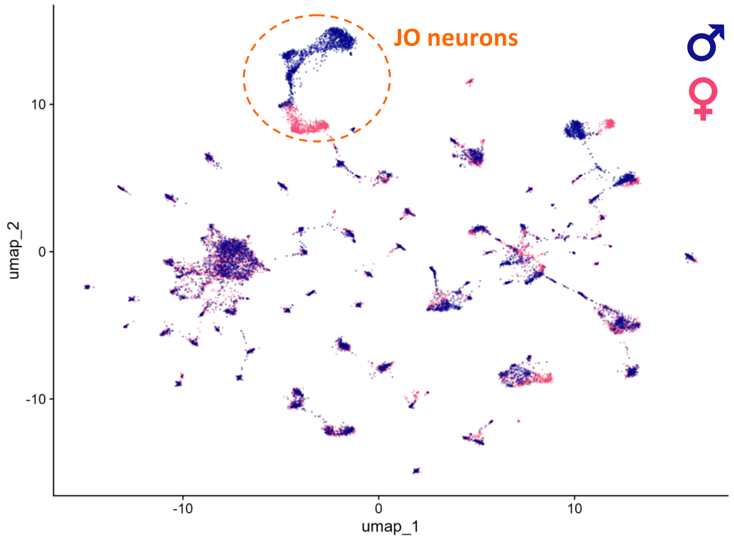

**B**

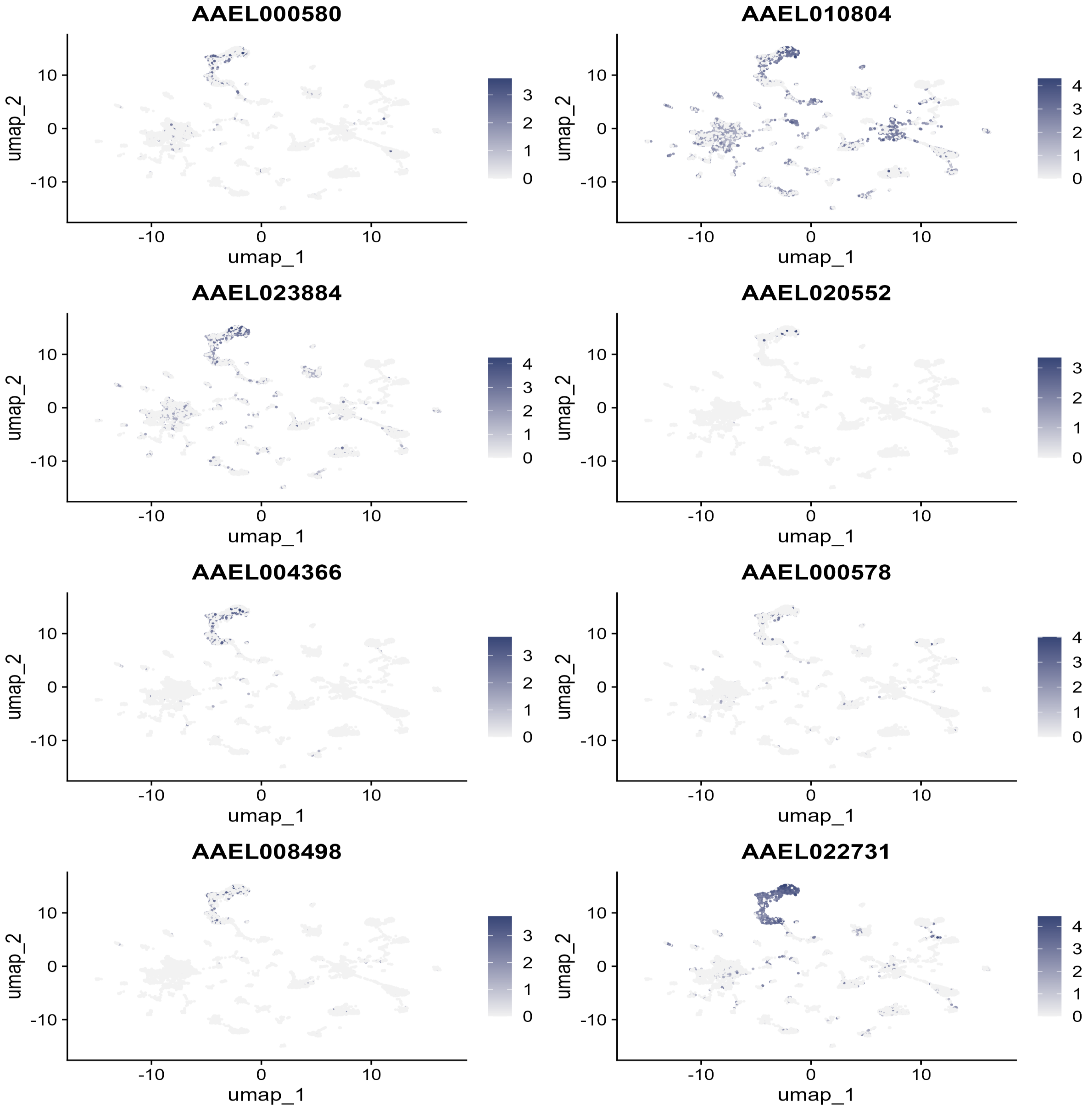

**Supplementary Figure 18: Expression levels of genes in $s_{242}$**

**A**

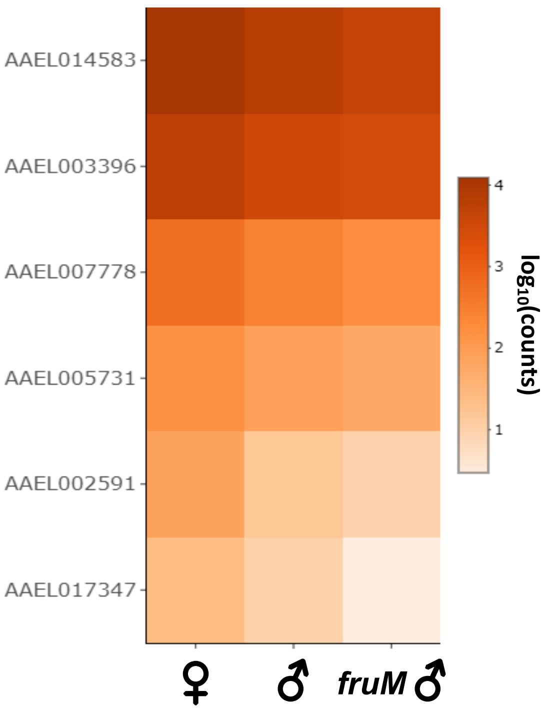

**Supplementary Figure 19: Identification of putative FruM-binding motifs**

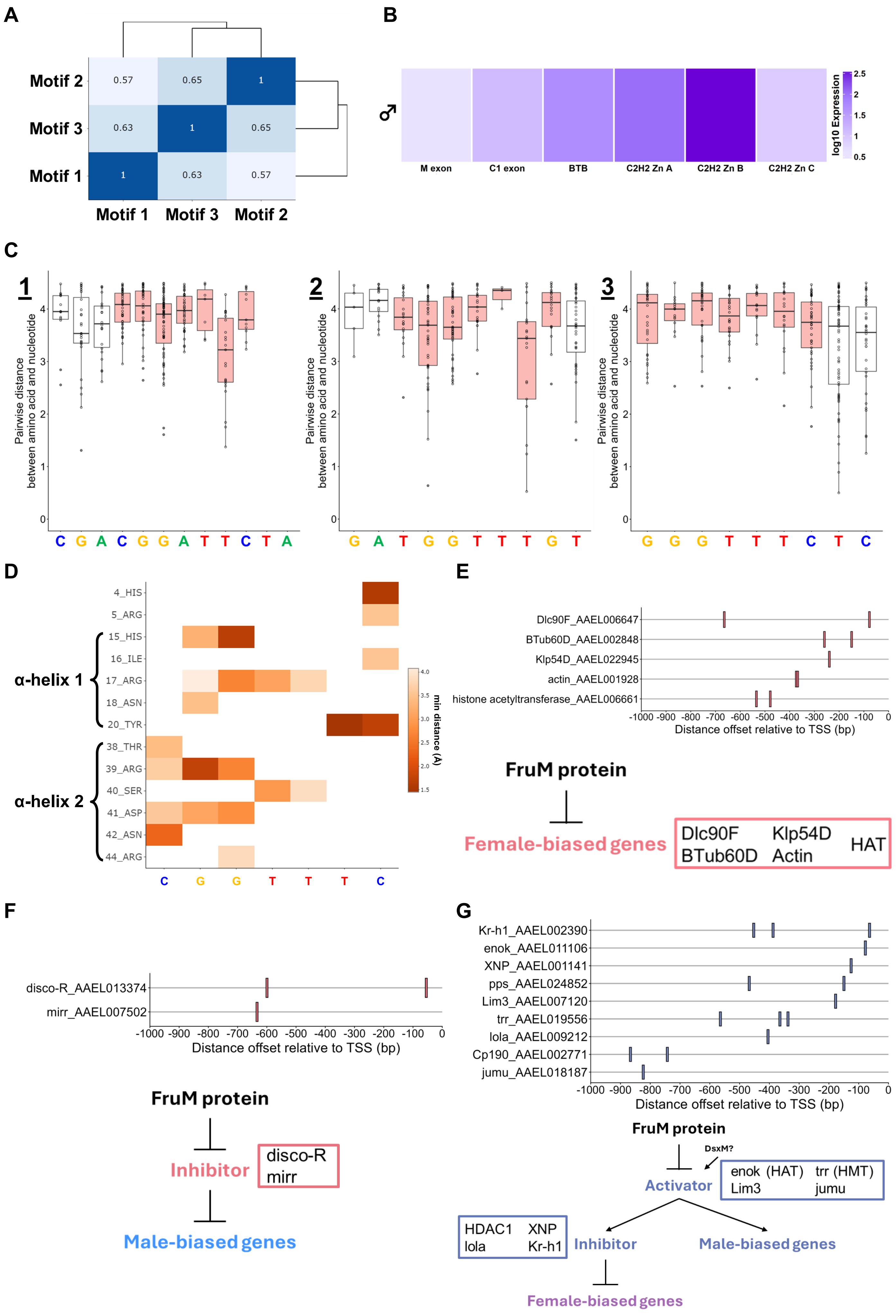

**Supplemental Information**

**Supplemental Figure Legends**

**Figure S1**. **Loss of FruM alters male locomotor activity**

**(A)** Locomotor Activity monitor diagram.

**(B)** Summed activity counts per hour for control females, control males and *fruM* males over three days of LD activity. Yellow shaded region represents lights on, grey shaded region represents lights off. Sample sizes: 40 control females; 49 control males; 58 *fruM* males.

**(C)** Summed beam breaks over 3 days for control female, control male and *fruM* male groups during ZT2-4 (left) and ZT11-12 (right). Individual points represent summed beam breaks for individual mosquitoes. Solid lines represent median and standard errors. n.s., $p>0.05$, *, $p<0.05$, **, $p<0.01$, ***, $p<0.001$, pairwise Wilcoxon tests with BH correction. See Tables S1 and S2 for p values. Sample sizes: 40 control females; 49 control males; 58 *fruM* males.

**Figure S2**. **Heatmaps of mosquito flight activity at dusk**

**(A)** Heatmaps of control female (top), control male (middle) and *fruM* male (bottom) pixel occupancy across three repeats. Each repeat contained 30 mosquitoes.

**Figure S3. Control female flight trajectories from BuzzWatch recordings**

**(A)** 30 longest trajectories from control female BuzzWatch recordings.

**Figure S4. Control male flight trajectories from BuzzWatch recordings**

**(A)** 30 longest trajectories from control male BuzzWatch recordings.

**Figure S5. *fruM* male flight trajectories from BuzzWatch recordings**

**(A)** 30 longest trajectories from *fruM* male BuzzWatch recordings.

**Figure S6. Loss of FruM abolishes male phonotaxis**

**(A)** Normalised frequency response profiles of control and *fruM* males to a range of pure tones (350-850 Hz in 25 Hz bins). Individual points represent individual repeat normalised frequency responses. Fig 1I shows same datasets over narrower frequency range (350-575Hz). Bars represent median normalised response per frequency across all repeats. Sample sizes: 2 control male cages; 5 *fruM* male cages.

**(B)** Normalized responses for $fruitless^{\Delta M}$/+ male cages to different frequencies of sound (350 – 850Hz in 25Hz steps). Individual points represent normalized responses for individual cages. Solid lines show average response between repeats. Responses normalized within a repeat. Sample size: 2 $fru^{\Delta M}$/+ male cages.

**Figure S7. *fru* expression in male JO and AMMC**

**(A)** Split channel images of *fru* expression in *fru*$^{\Delta M\text{-}tdTomato}$/+ male JO shown in Figure 2C. Scale bar, 100µm. Maximum intensity projections of JOs stained with anti- tdTomato (yellow) and anti-HRP (blue).

**(B)** *fru* expression in *fru*$^{\Delta M\text{-}tdTomato}$/+ male brain (left) focusing on Antennal Mechanosensory and Motor Center (AMMC) as well as whole brain image (right). Scale bar, 100µm. Maximum intensity projections of brains stained with anti- tdTomato (yellow) and anti-Brp (nc82, blue). Outlined regions in left images: red indicates AMMC, magenta indicates antennal lobe (AL) and green indicates JO neuron axon bundle. Brain images on left are from same brain with different layer coverage, image on right is from different brain.

**Figure S8. Loss of FruM alters male hearing function**

**(A)** Power spectral densities from harmonic oscillator fits to unstimulated free fluctuations of control female, control male and *fruM* male flagella in active and passive states. Thick lines represent fits generated based on median fit parameters. Grey lines represent passive system fits, colored lines represent active system fits. Sample sizes: 15 control females; 19 control males; 12 *fruM* males.

**(B)** Histogram of control female WBFs (red, Figure 1G) compared to control male (blue) and *fruM* male (orange) mechanical tuning frequencies in stimulated (sound, Figure 2E) and unstimulated (no sound, Figure S8A) states. Colored bars represent mechanical tuning frequencies across sound intensity gradient, from left (Sound/stimulated) to right (No sound/unstimulated).

**Figure S9. Loss of FruM alters localization of presynaptic terminals in male JOs**

**(A)** 3C11 staining within JO. Split channel images of JO IHC shown in Figure 3C. Scale bar, 100µm. Maximum intensity projections of JOs stained with anti-SYNORF1 (3C11, green) (top). Close-up views of type III (somata), type V (basal plate) and type VI (inner JO dendritic cilia) efferent presynaptic terminal sites taken from JO images shown in Figure 3C. Scale bar, 20µm. JOs stained with anti-SYNORF1 (3c11, green) and anti-HRP (magenta).

**(B)** nc46 staining within JO. Split channel images of JO IHC shown in Figure 3D. Scale bar, 100µm. Maximum intensity projections of JOs stained with anti- SAP47 (nc46, blue) (top). Close-up view of type III (somata), type V (basal plate) and type VI (inner JO dendritic cilia) efferent presynaptic terminal sites taken from JO images shown in Figure 3D. Scale bar, 20µm. JOs stained with anti-SAP47 (nc46, blue) and anti-HRP (magenta).

**Figure S10: Changes in mosquito hearing function following compound injection**

**(A)** Power spectral densities from harmonic oscillator fits to free fluctuations of control male flagella before (light blue) and after (dark blue) TeNT injection. Sample size: 3 control males.

**(B)** Maximum change in mechanical tuning frequency for vibrometry recordings taken after compound injection (Ringer or cAMP) relative to pre-injection baseline for control females, control males and *fruM* males. Individual points represent maximum changes in mechanical tuning after compound injection (Ringer or cAMP) compared to pre-injection baseline for individual mosquitoes. Solid lines show median and standard error. *, $p<0.05$; **, $p<0.01$; pairwise Wilcoxon test. See Tables S14 and S16 for maximum Δfrequency values and p values. Sample sizes for Ringer and cAMP injections: 7/10 control females; 8/8 control males; 8/8 *fruM* males.

**Figure S11: Gene ontology enrichment analysis**

**(A)** Network of GO enrichment terms identified as significantly enriched in genes significantly upregulated in *fruM* males compared to control males. Size and color of the GO term nodes represent the number of genes identified under each GO term. Thickness of the edges represents the number of shared genes between two GO term nodes. See Figure 4C.

**(B)** Network of GO enrichment terms identified as significantly enriched in genes significantly downregulated in *fruM* males compared to control males. Size and color of the GO term nodes represent the number of genes identified under each GO term. Thickness of the edges represents the number of shared genes between two GO term nodes. See Figure 4C.

**Figure S12: Expression levels of genes in $s_{142}$**

**(A)** Heatmap of expression levels of highlighted genes in $s_{142}$. See Figure 4D.

**(B)** Heatmap of expression levels of transcription factor genes in $s_{142}$.

**Figure S13: Expression of $s_{142}$ highlighted genes in Goldman et al, 2025 snRNA-seq database**

snRNA-seq database published in[1].

**(A)** UMAP of head nuclei of male and female *Ae. aegypti* colored by sex of head samples. Putative JO neurons, identified *via* expression of *inactive* (AAEL020482), are circled.

**(B)** UMAP of head nuclei of male and female *Ae. aegypti* colored by normalized expression of target gene in $s_{142}$. See Figure 4D.

**Figure S14: Expression levels of genes in $s_{175}$**

**(A)** Heatmap of expression levels of highlighted genes in $s_{175}$. See Figure 4E.

**Figure S15: Expression of $s_{175}$ highlighted genes in Goldman et al, 2025 snRNA-seq database.**

snRNA-seq database published in[1].

**(A)** UMAP of head nuclei of male and female *Ae. aegypti* colored by sex of head samples. Putative JO neurons, identified *via* expression of *inactive* (AAEL020482), are circled.

**(B)** UMAP of head nuclei of male and female *Ae. aegypti* colored by normalized expression of target gene in $s_{175}$. See Figure 4E.

**Figure S16: Expression levels of genes in $s_{257}$**

**(A)** Heatmap of expression levels of highlighted genes in $s_{257}$. See Figure 4F.

**(B)** Log2 fold change expression of gene of interest in $s_{257}$ that are upregulated in *fruM* males compared to control males and control males compared to control females. Transcriptional regulatory genes are highlighted.

**(C)** Heatmap of expression levels of transcriptional regulatory genes in $s_{257}$.

**Figure S17: Expression of $s_{257}$ highlighted genes in Goldman et al, 2025 snRNA-seq database.** snRNA-seq database published in[1].

**(A)** UMAP of head nuclei of male and female *Ae. aegypti* colored by sex of head samples. Putative JO neurons, identified *via* expression of *inactive* (AAEL020482), are circled.

**(B)** UMAP of head nuclei of male and female *Ae. aegypti* colored by normalized expression of target gene in $s_{257}$. See Figure 4F.

**Figure S18: Expression levels of genes in $s_{242}$**

**(A)** Heatmap of expression levels of highlighted genes in $s_{242}$. See Figure 4G.

**Figure S19: Identification of putative FruM-binding motifs**

**(A)** Pearson Correlation coefficients between members of FruM motif cluster. See Figure 5B.

**(B)** Heatmap of exon usage counts of different *fru* exons in male pedicel, from reanalysis of published wild-type adult male and female *Ae. aegypti* pedicel transcriptomes[2]. $C_2H_2$ Zn-A, Zn-B and Zn-C respectively refer to exons 37-39 of *fru* shown in Figure 2B.

**(C)** Pairwise distance analysis between Zn-B of FruM and FruM motif cluster members. All interactions < 4.5 Å between Zn-B and each nucleotide position plotted. Red shaded boxplots are nucleotide positions consistently showing interactions with Zn-B across all three models. See Figure 5B.

**(D)** Minimal distance interactions between indicated amino acids of native Zn-B of FruM and nucleotide positions of consensus FruM motif. Only interactions with minimal distance < 4.5 Å plotted. See Figure 5B for consensus model of interactions. Amino acid positions 15-20 form part of first alpha helix of Zn-B and amino acid positions 38-44 form part of second helix.

**(E)** Location of consensus FruM motif on 1kbp promoter sequence upstream of translational start site of genes highlighted in $s_{142}$, which could be female-biased genes under FruM's direct transcriptional inhibition. Only motifs showing conservation at positions 2, 3 and 5 with consensus motif analyzed. See Figures 4D, S12A and S13B.

**(F)** Location of consensus FruM motif on 1kbp promoter sequence upstream of translational start site of transcription factor genes in $s_{142}$. FruM could directly inhibit expression of these transcription factors, which could otherwise inhibit expression of male-biased genes in $s_{175}$. Only motifs showing conservation at positions 2, 3 and 5 with consensus motif analyzed. See Figures 4D and S12B.

**(G)** Location of consensus FruM motif on 1kbp promoter sequence upstream of translational start site of transcriptional regulatory genes in $s_{257}$. FruM could directly inhibit expression of transcriptional activators promoting expression of transcriptional inhibitors or male-biased genes in $s_{257}$. These transcriptional inhibitors could inhibit female-biased genes identified in $s_{242}$. Only motifs showing conservation at positions 2, 3 and 5 with consensus motif analyzed. See Figures 4F, 4G, S16B and S16C.

**Table S1. Statistical values for comparisons between groups for summed number of beam breaks during ZT2 - ZT4**.

Data collected from Locomotor Activity Monitor assays. Pairwise Wilcoxon tests with Benjamini Hochberg correction used for comparisons.

| Comparison groups | Adjusted p-values | W-values |
|---|---|---|
| Control male – Control female | 0.011 | 656.5 |
| *fruM* male – Control female | 0.610 | 1089.5 |
| *fruM* male – Control male | 0.011 | 991.5 |

**Table S2. Statistical values for comparisons between groups for summed number of beam breaks during ZT11 – ZT12.**

Data collected from Locomotor Activity Monitor assays. Pairwise Wilcoxon tests with Benjamini Hochberg correction used for comparisons.

| Comparison groups | Adjusted p-values | W-values |
|---|---|---|
| Control male – Control female | 0.331 | 1098.5 |
| *fruM* male – Control female | $2.15 \times 10^{-3}$ | 1601 |
| *fruM* male – Control male | $2.30 \times 10^{-5}$ | 2136.5 |

**Table S3: Statistical values for comparisons between groups for distance from centre of cage for BuzzWatch recordings**.

Data collected from BuzzWatch recordings. Pairwise Wilcoxon tests with Benjamini Hochberg correction used for comparisons.

| Comparison groups | Adjusted p-values | W-values |
|---|---|---|
| Control male – Control female | $8.76 \times 10^{-47}$ | 16021 |
| *fruM* male – Control female | 0.8713 | 29248 |
| *fruM* male – Control male | 0 | 76591 |

**Table S4: Statistical values for comparisons between groups for fly-by counts at dusk**. Data collected from Fly-by recordings. Pairwise t-tests with Benjamini Hochberg correction used for comparisons.

| Comparison groups | Adjusted p-values | t-values | Degrees of freedom |
|---|---|---|---|
| Control male – Control female | $3.58 \times 10^{-6}$ | 7.384 | 14.883 |
| *fruM* male – Control female | 0.247 | -1.2038 | 15.545 |
| *fruM* male – Control male | $3.58 \times 10^{-6}$ | -8.6574 | 11.568 |

**Table S5: Statistical values for comparisons between groups for Wing Beat Frequency.** Data collected from Fly-by recordings. Pairwise t-tests with Benjamini Hochberg correction used for comparisons.

| Comparison groups | Adjusted p-values | t-values | Degrees of freedom |
|---|---|---|---|
| Control male – Control female | $2.02 \times 10^{-21}$ | 51.162 | 19.091 |
| *fruM* male – Control female | $8.34 \times 10^{-15}$ | 34.548 | 14.036 |
| *fruM* male – Control male | 0.704 | 0.38674 | 15.929 |

**Table S6: Statistical values for comparisons between groups for mechanical tuning frequency during White Noise playback.**

Data collected using laser Doppler vibrometry during white noise stimulus playback. Pairwise t-tests with Benjamini Hochberg correction used for comparisons.

| Comparison groups | Adjusted p-values | t-values | Degrees of freedom |
|---|---|---|---|
| Control male – Control female | $3.36 \times 10^{-22}$ | 25.412 | 31.766 |
| *fruM* male – Control female | $2.05 \times 10^{-8}$ | 13.426 | 12.002 |
| *fruM* male – Control male | $2.99 \times 10^{-4}$ | 4.9411 | 12.547 |

**Table S7: Statistical values for comparisons between groups for mechanical tuning frequency in unstimulated, sedated state.**

Data collected using laser Doppler vibrometry from unstimulated, sedated mosquito ears. Pairwise t-tests with Benjamini Hochberg correction used for comparisons.

| Comparison groups | Adjusted p-values | t-values | Degrees of freedom |
|---|---|---|---|
| Control male – Control female | $1.98 \times 10^{-11}$ | 10.515 | 31.979 |
| *fruM* male – Control female | $1.66 \times 10^{-8}$ | 9.2989 | 19.896 |
| *fruM* male – Control male | 0.572 | 0.57353 | 22.225 |

**Table S8: Statistical values for comparisons between groups for mechanical tuning frequency during sweep stimulation.**

Data collected using combined vibrometry and electrophysiology paradigm utilising sweep stimuli. Pairwise t-tests with Benjamini Hochberg correction used for comparisons.

| Comparison groups | Adjusted p-values | t-values | Degrees of freedom |
|---|---|---|---|
| Control male – Control female | $6.69 \times 10^{-12}$ | 13.289 | 23.355 |
| *fruM* male – Control female | $7.21 \times 10^{-12}$ | 5.3299 | 25.405 |
| *fruM* male – Control male | $1.51 \times 10^{-5}$ | 17.923 | 16.065 |

**Table S9: Statistical values for comparisons between groups for electrical tuning frequency during sweep stimulation**.

Data collected using combined vibrometry and electrophysiology paradigm utilising sweep stimuli. Pairwise t-tests with Benjamini Hochberg correction used for comparisons.

| Comparison groups | Adjusted p-values | t-values | Degrees of freedom |
|---|---|---|---|
| Control male – Control female | $1.11 \times 10^{-9}$ | 10.05 | 22.854 |
| *fruM* male – Control female | $9.50 \times 10^{-19}$ | 23.248 | 26.648 |
| *fruM* male – Control male | $4.79 \times 10^{-5}$ | 4.9404 | 24.087 |

**Table S10: Statistical values for comparisons between groups for threshold minima electrical tuning frequency during pure tone stimulation**.

Data collected using combined vibrometry and electrophysiology paradigm utilising pure tone stimuli. Pairwise Wilcoxon tests with Benjamini Hochberg correction used for comparisons.

| Comparison groups | Adjusted p-values | W-values |
|---|---|---|
| Control male – Control female | $5.40 \times 10^{-8}$ | 7 |
| *fruM* male – Control female | $5.14 \times 10^{-9}$ | 0 |
| *fruM* male – Control male | $4.59 \times 10^{-3}$ | 59 |

**Table S11: Statistical values for comparisons between groups for minimum voltage required to stimulate a significant compound action potential during pure tone stimulation.** Data collected using combined vibrometry and electrophysiology paradigm utilising pure tone stimuli. Pairwise Wilcoxon tests with Benjamini Hochberg correction used for comparisons.

| Comparison groups | Adjusted p-values | W-values |
|---|---|---|
| Control male – Control female | $2.71 \times 10^{-4}$ | 248 |
| *fruM* male – Control female | 0.533 | 154 |
| *fruM* male – Control male | $1.11 \times 10^{-4}$ | 29 |

**Table S12: Statistical values for comparisons between groups for maximum action potential responses during pure tone stimulation**.

Data collected using combined vibrometry and electrophysiology paradigm utilising pure tone stimuli. Pairwise Wilcoxon tests with Benjamini Hochberg correction used for comparisons.

| Comparison groups | Adjusted p-values | W-values |
|---|---|---|
| Control male – Control female | $9.00 \times 10^{-9}$ | 3 |
| *fruM* male – Control female | 0.53 | 154 |
| *fruM* male – Control male | $9.00 \times 10^{-9}$ | 271 |

**Table S13: Statistical values for comparisons between groups for maximum action potential responses during step stimulation**.

Data collected using combined vibrometry and electrophysiology paradigm utilising pure tone stimuli. Pairwise t-tests with Benjamini Hochberg correction used for comparisons.

| Comparison groups | Adjusted p-values | t-values | Degrees of freedom |
|---|---|---|---|
| Control female – Control male | $1.03 \times 10^{-5}$ | -6.0034 | 20.245 |
| *fruM* male – Control female | $1.03 \times 10^{-5}$ | 6.5091 | 18.053 |
| *fruM* male – Control male | 0.828 | 0.21878 | 29.978 |

**Table S14: JO width measurements.**

All values provided as means ± standard deviations. Numbers in brackets refer to sample sizes for each experiment.

| Group | JO width (µm) |
|---|---|
| Control female | 138.44 ± 9.93 (n = 14) |
| Control male | 174.75 ± 8.74 (n = 14) |
| *fruM* male | 177.37 ± 11.10 (n = 13) |

**Table S15: Statistical values for comparisons between groups for JO width measurements.** Pairwise t-tests with Benjamini Hochberg correction used for comparisons.

| Comparison groups | Adjusted p-values | t-values | Degrees of freedom |
|---|---|---|---|
| Control female – Control male | $4.36 \times 10^{-10}$ | 10.269 | 25.583 |
| *fruM* male – Control female | $1.60 \times 10^{-9}$ | 9.577 | 24.157 |
| *fruM* male – Control male | 0.51 | 0.67769 | 22.814 |

**Table S16: Median values of maximum AUC ratio for Ringer and TeNT injection datasets (vibrometry).**

All values provided as medians ± standard errors. Numbers in brackets refer to sample sizes for each experiment.

| Group | Maximum AUC ratio after Ringer injection | Maximum AUC ratio after TeNT injection |
|---|---|---|
| Control female | 1.40 ± 0.10 (n = 3) | 1.23 ± 0.12 (n = 3) |
| Control male | 1.66 ± 0.36 (n = 3) | 83500 ± 19800 (n = 3) |
| *fruM* male | 1.39 ± 0.31 (n = 3) | 2.32 ± 0.66 (n = 4) |

**Table S17: Median values of maximum Δ frequency and AUC ratio for Ringer and cAMP injection datasets (vibrometry).**

All values provided as medians ± standard errors. Numbers in brackets refer to sample sizes for each experiment.

| Group | Maximum Δ frequency after Ringer injection (Hz) | Maximum Δ frequency after cAMP injection (Hz) | Maximum AUC ratio after Ringer injection | Maximum AUC ratio after cAMP injection |
|---|---|---|---|---|
| Control female | 11.06 ± 2.62 (n = 7) | 23.00 ± 9.88 (n = 10) | 1.23 ± 0.17 (n = 7) | 1.11 ± 0.19 (n = 10) |
| Control male | 31.35 ± 9.46 (n = 8) | 144.5 ± 12.03 (n = 8) | 1.72 ± 0.40 (n = 8) | 841.19 ± 3505.41 (n = 8) |
| *fruM* male | 11.64 ± 10.56 (n = 8) | 87.55 ± 16.95 (n = 8) | 1.90 ± 0.28 (n = 8) | 1.79 ± 0.31 (n = 8) |

**Table S18: Statistical values for maximum AUC ratio comparisons for Ringer and cAMP injection datasets.**

Data collected using laser Doppler vibrometry coupled with injections. Pairwise Wilcoxon tests used for comparisons.

| Group | Adjusted p-values | W-values |
|---|---|---|
| Control female | 0.860 | 41 |
| Control male | $1.55 \times 10^{-4}$ | 0 |
| *fruM* male | 0.442 | 40 |

**Table S19: Statistical values for maximum Δ frequency comparisons for Ringer and cAMP injection datasets.**

Data collected using laser Doppler vibrometry coupled with injections. Pairwise Wilcoxon tests used for comparisons.

| Group | Adjusted p-values | W-values |
|---|---|---|
| Control female | $9.67 \times 10^{-3}$ | 9 |
| Control male | $1.09 \times 10^{-3}$ | 3 |
| *fruM* male | $1.09 \times 10^{-3}$ | 3 |

**Table S20. (separate file)**

DESeq2 normalized counts for control female, control male and *fruM* male pedicel RNAsequencing data.

**Table S21. (separate file)**

DESeq2 output for comparisons between control female and control male pedicel RNAsequencing data.

**Table S22. (separate file)**

DESeq2 output for comparisons between *fruM* male and control female pedicel RNAsequencing data.

**Table S23. (separate file)**

DESeq2 output for comparisons between *fruM* male and control male pedicel RNAsequencing data.

**Table S24. (separate file)**

Output of HOMER FruM core consensus motif (7bp) scanning results for genes in $s_{142}$.

**Table S25. (separate file)**

Output of HOMER FruM core consensus motif (7bp) scanning results for genes in $s_{257}$.

**Table S26. (separate file)**

Output of HOMER FruM core consensus motif (7bp) scanning results for genes not differentially expressed between *fruM* male and control male pedicels (non-DEG group).

**Supplemental Files**

**Extended data Video 1. (separate file)**

Flight activity of groups of control female, control male and *fruM* male mosquitoes during ZT11.5 – ZT12. 30 mosquitoes per cage entrained for three days prior to recording.

**Extended data Video 2. (separate file)**

Representative video of tracking of *fruM* male flight trajectories using BuzzWatch system.

**Extended data Video 3. (separate file)**

Control male and *fruM* male responses to playback of 470 Hz tone. Left container houses 4 *fruM* males, right container 4 control males.
